# Divalent siRNA for prion disease

**DOI:** 10.1101/2024.12.05.627039

**Authors:** Juliana E Gentile, Taylor L Corridon, Fiona E Serack, Dimas Echeverria, Zachary C Kennedy, Corrie L Gallant-Behm, Matthew R Hassler, Garth A. Kinberger, Margaret N Kelemen, Nikita G Kamath, Yuan Lian, Katherine Y Gross, Rachael Miller, Kendrick DeSouza-Lenz, Michael Howard, Kenia Guzman, Nathan Chan, Vanessa Laversenne, Daniel T Curtis, Kevin Fettes, Marc Lemaitre, Aimee L Jackson, Ken Yamada, Julia F Alterman, Alissa A Coffey, Eric Vallabh Minikel, Anastasia Khvorova, Sonia M Vallabh

## Abstract

Prion protein (PrP) lowering is effective in animal models of prion disease and is being tested clinically in prion disease patients, but there remains a need for more potent PrP-lowering drug candidates. Inspired by the reported potency and duration of action of divalent short interfering RNA (siRNA), a new oligonucleotide drug modality for the central nervous system, we sought to discover and develop a new PrP-lowering drug candidate. Herein we identify a mouse *Prnp*-targeting divalent siRNA molecule, 1682-s4, that lowers PrP to 49% residual brain expression in wild-type mice, and, in the context of intracerebral infection with Rocky Mountain Laboratory (RML) prions, achieves a 2.7-fold increase in survival time with pre-symptomatic chronic treatment and 64% increase in survival time with a single dose after symptom onset. We describe the generation of two transgenic mouse lines, Tg25109 and Tg26372, expressing the full human *PRNP* gene and its non-coding sequence, and demonstrate their utility for in vivo discovery of potent human *PRNP*-targeting oligonucleotides. We discover siRNA sequence 2439 against human *PRNP* and compare its potency in different divalent siRNA chemical scaffolds. We determine that both the fixed UU tail and extended nucleic acid linkages of scaffold s4 contribute to superior potency compared to other scaffolds tested, offering 9.4 and 15.9 percentage points respectively of additional PrP knockdown. A single dose of 348 µg of 2439-s4 lowered whole brain hemisphere human PrP in transgenic mice to 17% residual after 30 days, while 52 µg lowered PrP to 49% residual. 1-2% of the dose of 2439-s4 delivered into cerebrospinal fluid is retained in the brain, and the median effective tissue concentration is estimated at 1.2 micrograms per gram of tissue. Good Laboratory Practices toxicology studies identified no significant liabilities, and the U.S. FDA has cleared an Investigational New Drug application to bring 2439-s4 into clinical trials.

## Introduction

Prion disease is a fatal, incurable neurodegenerative disease caused by misfolding of prion protein (PrP), encoded in humans by the gene *PRNP* (1). Convergent lines of evidence implicate PrP as the drug target in this disease (2), and indicate that lowering PrP should be both safe (3–9) and effective (10–15).The efficacy of PrP lowering at delaying onset and slowing progression of prion disease in animal models has been shown using antisense oligonucleotides (ASOs), zinc finger repressors, and a base editor (16–19). The therapeutic hypothesis of PrP lowering is now being tested clinically with an intrathecally administered PrP-lowering ASO, ION717, in symptomatic patients diagnosed with prion disease (NCT06153966).

We seek to augment the therapeutic pipeline for prion disease, for several reasons. Only 8-14% of drug candidates that enter Phase I ultimately reach approval (20–22), and while drug targets backed by human genetic evidence enjoy increased success rates (23), even these targets may take many drug candidates and many trials to yield a success (24). In mice, lowering by approximately half via heterozygous knockout or chronic early ASO treatment prolongs survival up to 3-fold, but all mice ultimately succumb to fatal prion disease (18), consistent with prion replication continuing, albeit at a reduced rate (24). Thus, to halt or indefinitely delay prion disease will require deeper than 50% target lowering.

Like ASOs, siRNAs are chemically modified oligonucleotide drugs that bind a target RNA through G·C and A·U base pairing, but whereas ASOs recruit RNase H1 to cleave the target RNA (25), siRNAs engage the RNA Induced Silencing Complex (RISC) to cleave their target (26). Like ASOs, siRNAs accumulate in endosomal depots (27), and their slow release from this compartment combined with their chemical stabilization allows them to provide months of pharmacologic effect following a single dose into cerebrospinal fluid (CSF). Divalent siRNA (28) is a novel siRNA architecture designed to enhance gene silencing within the central nervous system (CNS). It consists of two identical, fully chemically modified siRNA molecules connected by a linker, forming a larger molecule that distributes broadly in the brain following direct delivery into CSF and potently lowers its target RNA. The divalent scaffold has been shown to confer enhanced biodistribution and tissue uptake compared to a monovalent equivalent (28). Inspired by deep lowering of *HTT, APOE, SOD1*, and *KCNT1* in the rodent CNS (28–31)we set out to improve upon the divalent siRNA technology and to identify oligonucleotides against *PRNP* that knock down the target more than 50%. Divalent siRNA incorporates a variable number of phosphorothioate (PS) linkages at the 5′ and 3′ ends of each strand; PS is vital for cellular uptake and durability of oligonucleotide drugs but also mediates some toxicological properties, at least for single-stranded oligonucleotides (32, 33). We therefore also tested the hypothesis that the recently reported highly nuclease-resistant extended nucleic acid (exNA) nucleotide linkage (34) would permit us to reduce the number of PS linkages. In addition, because full complementarity to target RNA can result in RISC unloading or siRNA degradation (35, 36), we also sought to test the hypothesis that a fixed, non-complementary 3′ tail at the end of the antisense (AS) strand would improve potency.

Herein, we identify a tool compound against mouse *Prnp* and demonstrate that ∼50% PrP lowering with this new modality extends survival in prion-infected wild-type mice, replicating work with ASOs. We develop transgenic human *PRNP* mouse models and use them to identify a highly potent drug candidate against human *PRNP*, yielding as little as 17% residual PrP in whole brain hemisphere after a single dose. We demonstrate that both a fixed UU tail and the incorporation of exNA linkages contribute to the compound’s potency. We find activity out to 6 months post-dose and characterize the effect of repeat dosing on target engagement. Ultimately, we nominate a new drug candidate for prion disease.

## Methods

### Oligonucleotide synthesis

Oligonucleotides were synthesized by phosphoramidite solid-phase synthesis on automated synthesizer using a MerMade12 (Biosearch Technologies, Novato, CA), Dr Oligo 48 (Biolytic, Fremont, CA) or AKTA Oligopilot 100 (Cytiva, Marlborough, MA). 5ʹ-POM-vinyl phosphonate, 2ʹ-OMe-U CE-Phosphoramidite was used for the addition of 5′-Vinyl Phosphonate, 2ʹ-F, 2ʹ-OMe phosphoramidites with standard protecting groups were used to make the modified oligonucleotides. Phosphoramidites were dissolved at 0.1 M in anhydrous acetonitrile (ACN), with added anhydrous 15% dimethylformamide in the case of the 2′-OMe-Uridine amidite. 5-(Benzylthio)-1H-tetrazole (BTT) was used as the activator at 0.25 M. Detritylations was performed using 3% trichloroacetic acid in dichloromethane or Toluene. Capping reagents used were CAP A (20% N-methylimidazole in ACN) and CAP B (20% acetic anhydride and 30% 2,6-lutidine in ACN). Phosphite oxidation to convert to phosphate or phosphorothioate was performed with 0.05 M iodine in pyridine-water (9:1, v/v) or 0.1 M solution of 3-[(dimethylaminomethylene)amino]-3H-1,2,4-dithiazole-5-thione (DDTT) in pyridine, respectively. All reagents were purchased from Chemgenes, Wilmington, MA, phosphoramidites were purchased from Chemgenes and Hongene Biotech, Union City, CA. Oligonucleotides were grown on long-chain alkyl amine (LCAA) controlled pore glass (CPG) functionalized via succinyl linker with either Unylinker terminus for unconjugated oligonucleotides 500Å (Chemgenes), with cholesterol through a tetraethylene glycol linker 500 Å (Chemgenes), or with a di-trityl protected support separated by a tetraethylene glycol linker 1000 Å (Hongene Biotech) for divalent sense oligonucleotides.

For in cellulo experiments, oligonucleotides with or without cholesterol conjugate were cleaved and deprotected on-column with Ammonia gas (Airgas Specialty Gases). Briefly, columns were pre-wet with 100uL of water and immediately spun to remove the excess water. Columns were then placed in a reaction chamber (Biolytic) 90 min at 65°C. A modified on-column ethanol precipitation protocol was used for desalting and counterion exchange. Briefly, 1 mL of 0.1 M sodium acetate in 80% ethanol is flushed through the column, followed by a rinse with 1 mL 80% ethanol and finally after drying the excess ethanol, oligonucleotides were eluted with 600 µL of water in 96 deep well plates.

For in vivo experiments, 5′-(E)-Vinyl-phosphonate containing oligonucleotides were cleaved and deprotected with 3% diethylamine in ammonium hydroxide, for 20 h at 35℃ with agitation. Divalent oligonucleotides were cleaved and deprotected with 1:1 ammonium hydroxide and 40% aqueous monomethylamine, for 2 h at 25℃ with slight agitation. The controlled pore glass was subsequently filtered and rinsed with 30 mL of 5% ACN in water and dried overnight by centrifugal vacuum concentration. Purifications were performed on an Agilent 1290 Infinity II HPLC system using Source 15Q anion exchange resin (Cytiva). The loading solution was 20 mM sodium acetate in 10% ACN in water, and elution solution was the loading solution with 1M sodium bromide. Both oligonucleotide strands were eluted using a linear gradient from 30 to 70% in 40 min at 50°C. Peaks were monitored at 260 nm. Pure fractions were combined and desalted by size exclusion using Sephadex G-25 resins (Cytiva). Oligonucleotides were then lyophilized and resuspended in water.

Purity and identity of oligonucleotides were confirmed by IP-RP HPLC coupled to an Agilent 6530 Accurate-mass Q-TOF. LC parameters: buffer A: 100 mM 1,1,1,3,3,3-hexafluoroisopropanol (HFIP) (Oakwood Chemicals) and 9 mM triethylamine (TEA) (Fisher Scientific) in LC–MS grade water (Fisher Scientific); buffer B:100 mM HFIP and 9 mM TEA in LC–MS grade methanol (Fisher Scientific); column, Agilent AdvanceBio oligonucleotides C18; linear gradient 5–35% B 5 min was used for unconjugated and divalent oligonucleotides; linear gradient 25–80% B 5 min was used for cholesterol conjugated oligonucleotides; temperature, 60°C; flow rate, 0.85 ml/min. Peaks were monitored at 260nm. MS parameters: Source, electrospray ionization; ion polarity, negative mode; range, 100–3,200 m/z; scan rate, 2 spectra/s; capillary voltage, 4,000; fragmentor, 200 V; gas temp, 325°C.

### Initial screen for human and mouse siRNA sequences

Screening of siRNA sequences in cellulo utilized gymnotic uptake of monovalent cholesterol tetraethylene glycol (Chol-TEG) conjugated siRNAs, as cholesterol conjugates have demonstrated efficient gymnotic uptake into cultured cells and utility as tools for screening (37). The initial screening for potent siRNA sequences was conducted at UMass Chan, including both the synthesis of siRNAs in the monovalent **Chol-TEG s1** scaffold (Figure S1) and the cell culture screening experiments. Sequences were bioinformatically nominated by a published algorithm (38). Sense and antisense strands were annealed together at 95°C for 10 minutes and then cooled to room temperature. Screens were performed in N2a cells (mouse; ATCC No. CCL-131) or A549 cells (human; ATCC No. CCL-185) which were previously reported to have robust PrP expression (39). Cells were seeded into triplicate wells with growth media containing 1.5 µM of compound. After 72 hours, cell lysates were harvested. Mouse *Prnp* / human *PRNP* RNA were quantified with Quantigene assays (QGS-1000 SB-3030881 and SA-3002866 respectively), and as housekeeping controls, *Hprt/HPRT* RNA were quantified (assays SB-15463 and SA-10030 respectively) using the QuantigeneTM 2.0 branched DNA assay (Invitrogen). The ratio of *Prnp/PRNP* to *Hprt/HPRT* was normalized to the mean of expected non-targeting sequences (meaning, data for human cells were normalized to the mean of mouse-only sequences, while data for mouse cells were normalized to the mean of human-only sequences) to obtain an estimate of residual target expression after siRNA treatment. Highly active compounds found in the screen were then assayed in triplicate across 7 doses (23 nM – 1.5 μM) to calculate half-maximal inhibitory concentration (IC_50_). The numbering of human siRNA sequences used herein is relative to the transcription start site of a now-outdated RefSeq transcript NM_000311.4, though we note that human brain expression data support exclusive use of canonical Ensembl transcript ENST00000379440.9 which begins 362 bases further downstream (40).

### Expanded screen for potent human siRNA sequences

Further screening for potent human siRNA sequences was conducted with siRNAs synthesized in the monovalent **Chol-TEG s2** scaffold at Atalanta Therapeutics and tested in cellulo at the Broad Institute. Oligonucleotides were synthesized with standard solid-support phosphoramidite chemistry using a Dr. Oligo synthesizer. For single point primary screens siRNAs were diluted to twice the desired final concentration in optiMEM (Gibco cat no: 31985070) and for IC50 determination a 9-point 1:3 serial dilution series was used. U-251 MG glioblastoma cells (Sigma Aldrich cat no: 09063001) growing in a T75 flask were washed with PBS, trypsinized then quenched with growth media: optiMEM, 10% FBS, 1% NEAA (Gibco cat no: 11140050), 1% GlutaMAX (Gibco cat no: 35050061), 1% pen/strep (Gibco cat no: 15140122). Cells were pelleted at 1000 rpm for 5 minutes, growth media was aspirated then cells were resuspended in media containing 6% FBS without pen/strep. Sterile PBS was placed in outer wells of a 96-well plate to limit evaporation, while 8,000 cells per well were plated on the inner 60 wells. 50 µL of prepared cells was added to every well. 50 µL of optiMEM with 2x siRNA was added to the treated wells. Every plate contained 6-wells of “untreated cells” (50 µL optiMEM with no additives) for assay normalization purposes. Each condition was tested in biological triplicate (three cell culture wells) and each biological replicate was analyzed by qPCR in technical duplicate. After 72 hrs, any wells with cell death or rounded cells were noted then cells were lysed using the Cells-to-CT 1-step TaqMan Kit (Invitrogen cat no: A25602) according to the manufacturer protocol, with slight deviations. Media was aspirated from all wells then cells were washed with 200 µL 4°C sterile PBS. PBS was aspirated from every well before adding lysis solution containing DNase. Plate was placed on a shaker for 5 minutes then stop solution was added to every well. Cells were placed back on a shaker for 3 minutes then cell lysate was stored. RT-PCR samples were prepared using 2 µL or cell lysate, Taqman 1-Step qRT-PCR master mix and Taqman gene expression assays for human *TBP* (Invitrogen, cat no: Hs00427620_m1) and human *PRNP* (Invitrogen, cat no: Hs00175591_m1) in a 20 µL reaction volume. Samples were run on a QuantStudio 7 Flex system (Applied Biosystems) in a MicroAmp Optical 384-well reaction plate (Applied Biosystems, cat no: 4309849) using the following cycling conditions: reverse transcription 50°C, 5 min; reverse transcription inactivation/initial denaturation 95°C, 20 s; amplification 95°C, 3 s, 60°C, 30 s, 40 cycles. Each biological sample was run in duplicate, and the level of all targets were determined by ΔΔCt whereby results were first normalized to the housekeeping gene TBP and then to the untreated samples.

### In silico off-targets analysis

We opted not to pursue a comprehensive transcriptomic evaluation of off-targets in cell lines, because complementarity of the seed region (bases 2-8) alone can drive off-target profiles of siRNA in cultured cells (41) whereas one report indicates that seed complementarity may not be a major contributor to off-target activity for divalent siRNA in vivo (28). An in silico strategy for assessing off-target risks was considered adequate by FDA for IND filing. The reverse complement of bases 2-17 of the 2439 sequence — CACTTTGTGAGTATTC — was searched in NCBI Nucleotide BLAST (https://blast.ncbi.nlm.nih.gov/Blast.cgi) with the following settings. Database: Standard databases (nr etc.): > RefSeq Select RNA sequences (refseq_select). Organism: Homo sapiens (taxid:9606). Optimize for: Highly similar sequences (megablast). Max target sequences: 5,000. Match/mismatch scores: 1,-1. All other settings default. The top 100 hits from BLAST were downloaded as an XML file and parsed into tabular form using a Python script generated by ChatGPT. The table was then filtered using a custom R script to include only matches where the search string was antisense to the target, consistent with the siRNA mechanism, and was sorted by number of mismatches.

### Transgenic mouse generation

Transgenesis was performed by Cyagen (Santa Clara, CA) at its site in Taicang, Jiangsu, China. Searching NCBI CloneDB nominated bacterial artificial chromosome (BAC) RP11-715K24 as overlapping human PRNP. After sequence confirmation, *PRNP* and flanking sequence were cloned into the pStart-K plasmid by gap-repair cloning, with the p15A origin of replication and kanamycin resistance cassette located downstream of the gene, yielding a 48kb plasmid whose sequence is provided in this study’s online git repository. The plasmid was linearized with restriction enzyme NruI and microinjected into fertilized C57BL/6 eggs. PCR screening identified four founder pups, of which animals 26372 (male) and 25109 (female) were successfully bred. Transgenic animals were backcrossed to wild-type C57BL/6N animals until generation F5, then crossed to ZH3 PrP knockout mice (42) (on a C57BL/6J background) housed at McLaughlin Research Institute, with rederivation to remove opportunistic pathogens performed by Charles River Labs. Mouse lines have been deposited with Mutant Mouse Resource & Research Centers (MMRRC) for both academic and for-profit groups to utilize freely (accession numbers Tg26372: RRID:MMRRC_075939-UCD; Tg25109: RRID:MMRRC_075940-UCD).

### Transgene characterization

Transgene mapping was performed by Taconic and Cergentis using targeted locus amplification (TLA) (43) on spleens from 9-week-old males of generation F3 (Tg26372) and F6 (Tg25109). Copy number was estimated based on the number of transgene-transgene fusion reads and the ratio of transgene coverage to flanking region coverage. Full transgene characterization reports are provided in this study’s online git repository. Sequencing of the Tg26372 mouse utilized custom capture probes by Twist Bioscience targeting 152kb of human sequence surrounding PRNP as previously described (44). Zygosity-aware genotyping was performed by Transnetyx.

### Dose levels of siRNA

Drugs were formulated to target doses in terms of nanomoles (nmol) total bilateral dose, with the molarity referring to the full divalent siRNA molecule and not the monovalent equivalents. We targeted dose levels of 10, 5, 1.5, 1, or 0.2 nmol, based on siRNA concentrations determined by UV absorbance on NanoDrop. At UMass Chan (s1 and s4 scaffolds), concentration determinations initially used the theoretical molar extinction coefficients (MECs) determined by the base composition method, in which the extinction coefficients of each base are summed and multiplied by 0.9 to account for base stacking. For instance, for our drug candidate 2439-s4, we formulated drug for mouse studies based on the theoretical MEC of 766,260 L⋅mol^−1^⋅cm^−1^ (twice the sum of the antisense and sense strand MECs). The theoretical base composition MECs, however, do not account for hypochromicity — oligonucleotides absorb less UV light when duplexed. At Atalanta Therapeutics (s2 and s3 scaffolds), the nearest-neighbors (NN) was used, which attempts to account for the hyperchromicity in a sequence-specific manner. For 2439-s2, this yielded a theoretical MEC of 656,658 L⋅mol^−1^⋅cm^−1^. Later in the development program the MEC of 2439-s4 was determined empirically to be 549,107 L⋅mol^−1^⋅cm^−1^. This indicated that all doses of compounds synthesized at UMass Chan had been 1.395 times higher than believed at the time. Thus, for instance, studies of 2439-s4 intended to use a 10 nmol dose were in fact performed at a 13.95 nmol dose level. The 13.95 nmol dose, multiplied by the 2439-s4 molecular weight of 24,952 Da, corresponds to 348 µg. Empirical MECs were not determined for 2439-s2 nor any other compounds tested here, however, in reporting doses in this manuscript, we have applied this same correction factor across all compounds described herein. We originally sought to evaluate dose levels of 10, 5, 1.5, 1, or 0.2 nmol, based on the reported tolerability of divalent siRNAs up to 10 nmol (28). Based on the empirical MEC of 2439-s4, we estimate that those dose levels correspond respectively to 348, 174, 52, 35, and 7 µg, which assumes a 766,260/549,107 = 39.5% difference between empirical and uncorrected extinction coefficients as determined for 2439-s4. We note, however, that the dose of 2439-s2 and other s2 and s3 scaffold compounds that we administered had already accounted for a smaller, theoretically calculated hypochromicity adjustment, thus, our approach may be an overcorrection. If the hypochromicity of the compounds synthesized at Atalanta is similar to that of 2439-s4, then the doses of Atalanta compounds tested herein may be 14% lower than those of UMass Chan compounds (for example, 298 µg versus 348 µg).

### Animal studies

All animal studies were conducted under Broad Institute IACUC protocol 0162-05-17. Transgenic mice (see above) were bred at the Broad Institute. Wild-type C57BL/6N mice were purchased from Charles River Laboratories.

### Administration of divalent siRNA to mice

For in vivo use, siRNAs were synthesized in divalent format by UMass Medical School or Atalanta Therapeutics, see detailed methods above. Atalanta compounds were formulated in 1X phosphate-buffered saline (PBS) without ionic conditioning. At UMass Chan, to prevent neurotoxicity due to divalent cation imbalance (45–47), siRNAs were formulated with divalent cations: stock solutions of 1 mM divalent siRNA (2 mM monovalent equivalents) were prepared with 2 mM MgCl_2_, 14 mM CaCl_2_, 8 mM HEPES, 20 mM D-glucose, 5 mM KCl, and 137 mM NaCl. Dilutions of this stock were prepared in artificial CSF containing 137 mM NaCl, 5 mM KCl, 20 mM D-glucose, and 8 mM HEPES. siRNAs were delivered to mice via bilateral intracerebroventricular (ICV) injection, 5 µL per side for a total of 10 µL injection volume. The ICV procedure was slightly modified from that described previously for antisense oligonucleotides (18). In all in vivo target engagement studies, mice were 4-18 weeks old at the time of first ICV injection. In the first 4 studies of in vivo target engagement, animals were anesthetized with 1.2% tribromoethanol, injected i.p. with 0.23 mL per 10 g of body weight using an insulin syringe (BD 329410). Tribromoethanol was prepared freshly each week according to a published protocol (48), passed through a 0.22 µm filter, handled under sterile conditions and stored at 4°C in the dark until use, discarding if stored past 2 weeks. In the remaining 12 studies of in vivo target engagement and in the survival studies in the prion disease model, we used 3% isoflurane inhalation anesthesia. Anesthetized animals were immobilized in a stereotactic apparatus (SAS-4100, ASI Instruments) with 18° ear bars and the nose bar set to -8 mm. Heads were shaved and scalps swabbed with povidone/iodine and alcohol swabs, a 1 cm incision was made along the midline and the periosteum was scrubbed with a sterile cotton-tipped applicator to reveal the bregma landmark. Microliter syringes with either 26 G or 22 G needles (Hamilton company model 701, point style 2, No. 80300 or 80308 respectively) were filled with 10 µL of formulated drug or vehicle. From bregma, the needle was moved 0.3 mm rostral, 1.0 mm right or left, then down until it touched the skull and then 3.5 mm ventral. 5 µL was ejected gradually over ∼10 seconds, then after a 1 minute pause the needle was backed out while maintaining downward pressure on the skull with a cotton-tipped applicator. This procedure was performed first on the right and then on the left. After both injections, incisions were closed with either wound clips (Braintree Scientific cat no: RF7) or sutured with a single horizontal mattress stitch (Ethicon 661H). In target engagement studies, animals were generally harvested after 4 weeks, and a minimum of 3 weeks, in-life, to permit a sufficient number of PrP half-lives to observe lowering at the protein level (49). Because ASOs were re-dosed every 90 days in chronic treatment studies and both our and other data (28) have shown a long durability of divalent siRNA activity in the brain, we chose a re-dose interval of 120 days for our survival studies.

### Prion infection

Prion inoculations were performed as described previously (17, 18). To prepare the challenge agent, brains of terminally prion-sick mice infected with the Rocky Mountain Laboratories (RML) prion strain (50) were frozen, homogenized at 10% wt/vol in phosphate-buffered saline (PBS; Gibco 14190) using 1.4 mm zirconium oxide beads in 7 mL screw-cap tubes (CK14 soft tissue homogenizing kit, Precellys KT039611307.7) by means of 3x 40-second pulses on high in a Minilys homogenizer (Bertin EQ06404-200-RD000.0). 10% homogenate was then diluted 1:10 (vol/vol) to yield a 1% homogenate, irradiated with 7 kGy of X-rays on dry ice, extruded through finer and finer blunt-end needles (Sai infusion B18, B21, B24, B27, B30), and injected into sterile amber glass vials (MedLabSupply) and frozen. On the day of inoculation, vials were thawed and for each animal, 30 µL was drawn into a disposable insulin syringe with a 31 G 6 mm needle (BD SafetyGlide 328449). 7-week old C57BL/6N animals (Charles River) were placed under 3.0% isoflurane inhalation anesthesia, received meloxicam (5 mg/kg) for analgesia (one dose prophylactically and post-operative doses on following days), and heads were swabbed with povidone/iodine and alcohol swabs. The needle was then freehand inserted through the skull between the right ear and midline. After three seconds, the needle was withdrawn and animals were returned to home cages.

### Animal monitoring

All animals undergoing ICV drug administration received post-operative monitoring daily for 4 days to surveil recovery and wound closure. In target engagement studies in non-prion animals, weights were generally collected prior to dosing and at 1-week intervals thereafter, although staffing constraints led to weights not being consistently collected in a subset of studies. Prion-infected animals had baseline body weights taken at 16 weeks of age (corresponding to 60 days-post inoculation, dpi) and weekly thereafter until 120 dpi, after which weights were taken thrice weekly. On the same monitoring schedule, we also collected behavioral scores and nest scores as described (18). Behavioral scores were rated as 0 = absent, 1 = present for 8 symptoms: scruff / poor grooming, poor body condition, reduced activity, hunched posture, irregular gait/hindlimb weakness, tremor, blank stare, and difficulty righting. Nest scores were assigned for both cotton square nestlets (Ancare) and Enviro-dri® packed paper (Shepherd): 0 = unused; 1 = used/pulled apart, but flat; 2 = pulled into a three-dimensional structure; 0.5 and 1.5 were permitted intermediate scores. Animals were euthanized by CO2 inhalation (Euthanex) when they reached -20% weight loss relative to individual baseline, or were deemed moribund, defined as unable to reach food or water. All monitoring was conducted, and endpoint decisions taken, by veterinary technicians blinded to treatment group (PBS vs. non-targeting vs. active compound), although in studies with no injection controls, the lack of drug treatment in the control group could be inferred from the absence of surgery cards. As in our previous work, survival curves include deaths of all causes — animals are excluded only in cases of 1) death due to surgical complications on the day of surgery, 2) death prior to treatment group assignment, or 3) experimental error (for instance, wrong dose administered).

### Tissue processing, PrP quantification, and RNA analysis

All quantification of PrP protein was performed on whole mouse brain hemispheres including cerebellum. Our in-house PrP ELISA has been previously described (51) and is summarized briefly as follows. Whole hemispheres were frozen on dry ice and later homogenized at 10% wt/vol in 0.2% CHAPS. The assay uses antibodies EP1802Y (Abcam, ab52604) for capture and in-house biotinylated 8H4 (Abcam ab61409) for detection, followed by streptavidin-HRP (Thermo Fisher Scientific, 21130) and TMB (Cell Signaling, 7004P4). Our calibrator curve from 5 ng/mL to 0.05 ng/mL utilized recombinant full-length mouse PrP (MoPrP23-231) expressed in *E. coli* and purified in-house (52). We have previously shown (51) that this ELISA assay has indistinguishable reactivity for human and mouse PrP. Wild-type and Tg25109 brains were run at a 1:200 final dilution (10% homogenate diluted 1:20), while Tg26372 brains, because they overexpress PrP, were run at a 1:400 dilution. Each plate included high (WT), mid (het KO), and low (10% WT / 90% KO mix) brain homogenates used as QCs. To control against plate-to-plate variability, whenever one experiment produced more samples than could be run on one plate, we split every treatment group equally across 2 or more plates and normalized each sample to the mean of the PBS or no injection controls on its same plate. All results are expressed as residual PrP, a percentage of the control level. Across all experiments described here, the mid and low QCs usually read out at greater than the expected values of 50% and 10% residual PrP respectively, suggesting that the assay often overestimates residual PrP; for instance, the low QC, designed to mimic 10% residual PrP, averaged 17% residual PrP across all ELISA plates run in this study. Summary statistics on all ELISA plates are available in Figure S6. For RNA quantification, fresh brain hemispheres were placed in RNAlater (Sigma cat no: R0901) at 4°C before dissecting brain regions as described (51). RNA was extracted from brain tissue using a Qiagen RNeasy Lipid Tissue mini Kit (cat no: 74804) with few deviations from the manufacturer protocol. Whole hemispheres were homogenized at 10% wt/vol in QIAzol then 100 mg of tissue was applied to column. On-column DNase digestion was not performed and finally RNA was eluted into 40 µL RNase-free water. RNA yield and purity was checked by nanodrop. RT-qPCR was performed as described above using Taqman gene expression assays for mouse *Tbp* (Invitrogen, cat no: Mm00446971_m1) and human *PRNP* (Invitrogen, cat no: Hs00175591_m1) for transgenic mouse models or mouse Prnp (Invitrogen, cat no: Mm00448389_m1) for wild-type C57BL/6N mice. PBS or no injection animals were used as the control group.

### Pharmacokinetic analysis

Pharmacokinetic (PK; drug concentration in tissue and biofluid) measurements were performed at Axolabs GmbH (Kulmbach, Germany). A peptide nucleic acid (PNA) probe was designed to be complementary to the antisense strand of sequence 2439: (C term) Atto425-OO-cdctttgtgdgtd (N-term). This probe was allowed to hybridize to drug present in tissue homogenates, plasma, or CSF, and then run on anion exchange (AEX) high performance liquid chromatography (HPLC). The assay principle of PNA-HPLC is that the area under the curve of the fluorescent PNA-antisense duplex in the HPLC elution is integrated to derive the drug concentration (53). The lower limit of quantification (LLOQ) was determined to be 1 ng/mL in plasma and CSF, and 10 ng/mL in tissue homogenate. Because we did not have pharmacokinetic measurements in mouse CSF, we were unable to perform the complex multi-compartment pharmacology models reported for certain other oligonucleotides (54). Instead, simple 4-point Hill slope models were fit to model the effect of either drug dose administered or drug accumulation in tissue (independent variable) on target knockdown (dependent variable), see “Statistics, source code, and data availability” for details.

### Investigational New Drug (IND)-enabling studies

56.5 g (gross weight) of lyophilized 2439-s4 (Drug Substance) was produced at at Hongene Bioengineering (Shanghai, China) via manufacture process transferred from UMass (see above) and scaled up using starting materials manufactured by Hongene and equivalent reagents by Hongene or qualified local suppliers. The release testing methods had been developed and qualified at Hongene, and the manufacturing and release confirmed to Good Manufacturing Practices (GMP) for clinical products. The purity of drug substance was 96.4% by non-denaturing size exclusion chromatography. GMP material was used both for production of sterile injectable formulation (Drug Product) by Argonaut Manufacturing Services (Carlsbad, California, USA), and for toxicology studies.

Good Laboratory Practices (GLP) toxicology studies by single lumbar intrathecal stick (no catheter) in beagle dogs and Sprague-Dawley rats were performed by Amplify-Bio (West Jefferson, Ohio, USA). Four dose levels were tested in each species with takedowns scheduled for 24h (core cohort) and 28d post-dose (recovery cohort) (see Results for full study designs). In-life assessments for rats included: moribundity and mortality, cage-side clinical observations, weekly body weights, food consumption, and ophthalmic exams. In-life assessments for dogs included: cage-side clinical observations, moribundity and mortality, weekly body weights, food consumption, noninvasive homecage neurobehavioral assessment, ophthalmic exams, electrocardiogram (ECG), noninvasive blood pressure, and respiratory rate. Blood samples were collected from pre-dose, 0.5, 4, 8, 12, 24, 48, and 72 hours post dose for toxicokinetic (TK) evaluation. Blood was collected at necropsy for hematology, serum chemistry, and coagulation evaluations. Urine was collected for urinalysis. CSF was collected for biodistribution. Tissue was collected for biodistribution and histopathology. Tissue was processed to slides and stained with hematoxylin and eosin for histopathologic examination by a board-certified veterinary pathologist. Drug concentrations in blood, CSF, and tissue were determined by Axolabs (Kulmbach, Germany) using peptide nucleic acid (PNA) hybridization and high performance liquid chromatography (see above).

GLP in vitro Ames and micronucleus genotoxicity tests were performed by ITR Laboratories (Montreal, Canada). The cytogenetic potential of 2439-s4 was tested using the in vitro micronucleus test with Chinese hamster ovary derived CHO-K1 cells with 4h incubation with rat liver S9 microsomal fraction (metabolic activation present) or with 4h or 26h incubation without S9 (metabolic activation absent). The mutagenic potential of 2439-s4 was tested using the Ames Test in four *Salmonella* strains and one *E. coli* strain with or without S9, a rat liver extract containing microsomal enzymes which mimics metabolism. Seven doses were tested in multiple strains (see Results).

Non-GLP drug-drug interaction studies were performed by BioIVT (Kansas City, Kansas, USA). Cytochrome P450 (Cyp) inhibition studies were performed in human hepatocytes with 0min or 30min pretreatment with 2439-s4. Cyp enzymes tested were: CYP1A2, CYP2B6, CYP2C9, CYP2C19, CYP2D6, CYP3A4/5 (probe substrates: midazolam and testosterone), CYP2C8. Cyp induction studies were performed in human hepatocytes with 72h incubation. Expression of Cyp enzymes was measured using qRT-PCR. Cyp enzymes tested were: CYP1A2, CYP2B6, CYP3A4. Transporter inhibition studies tested the effect of 2439-s4 on transport of a substrate across a membrane using Caco-2 cells for permeability-glycoprotein (P-gp), MDCKII cells for breast cancer resistance protein (BCRP), or HEK293 cells expressing the transporter of interest for OATP1B1, OATP1B3, OAT1, OAT3, OCT2, MATE1, and MATE2-K. MDCKII cells expressing the transporter of interest were used to test whether 2439-s4 is a substrate of P-gp or BCRP.

Further methodological details are not elaborated here, as final study reports and IND documents have been provided publicly (see below).

### Statistics, source code, and data availability

All analysis was conducted using custom scripts in R 4.2.0. Percent changes in mouse weights relative to baseline were compared using 2-sided T tests. Survival was assessed using log-rank test. Dose-response curves were fit using the drc package (50) in R, with a 4-point Hill slope model fixing the infinite-dose asymptote (c) and zero-dose asymptote (d) at 0% and 100% respectively. The impact of scaffold and fixed tail was characterized using a linear model with formula residual ∼ scaffold + log(dose) + region. Inflammatory marker responses were assessed using Dunnett’s test to compare each treated group to the PBS or untreated group. Raw individual-level animal data, source code sufficient to reproduce all analyses herein, and the text our Investigational New Drug application and regulatory interactions with U.S. FDA are available in this study’s online git repository: https://github.com/ericminikel/divalent

## Results

### Divalent siRNA chemical scaffolds

We employed several distinct chemical scaffolds to test divalent siRNAs in vivo (Figure 1). The previously reported **s1** scaffold (29) and the **s2** variant with one additional 2ʹ-OMe modification possess 7 and 5 phosphorothioate (PS) linkages at the 3′ end of the antisense (AS) strand respectively (Figure 1). In the **s3** scaffold, the AS 3′ end is reduced to just 2 PS linkages, and in **s4**, these two 3′ nucleotides are further stabilized with the extended nucleic acid (exNA) modification and are always uracil (U) (34).

**Figure 1.**
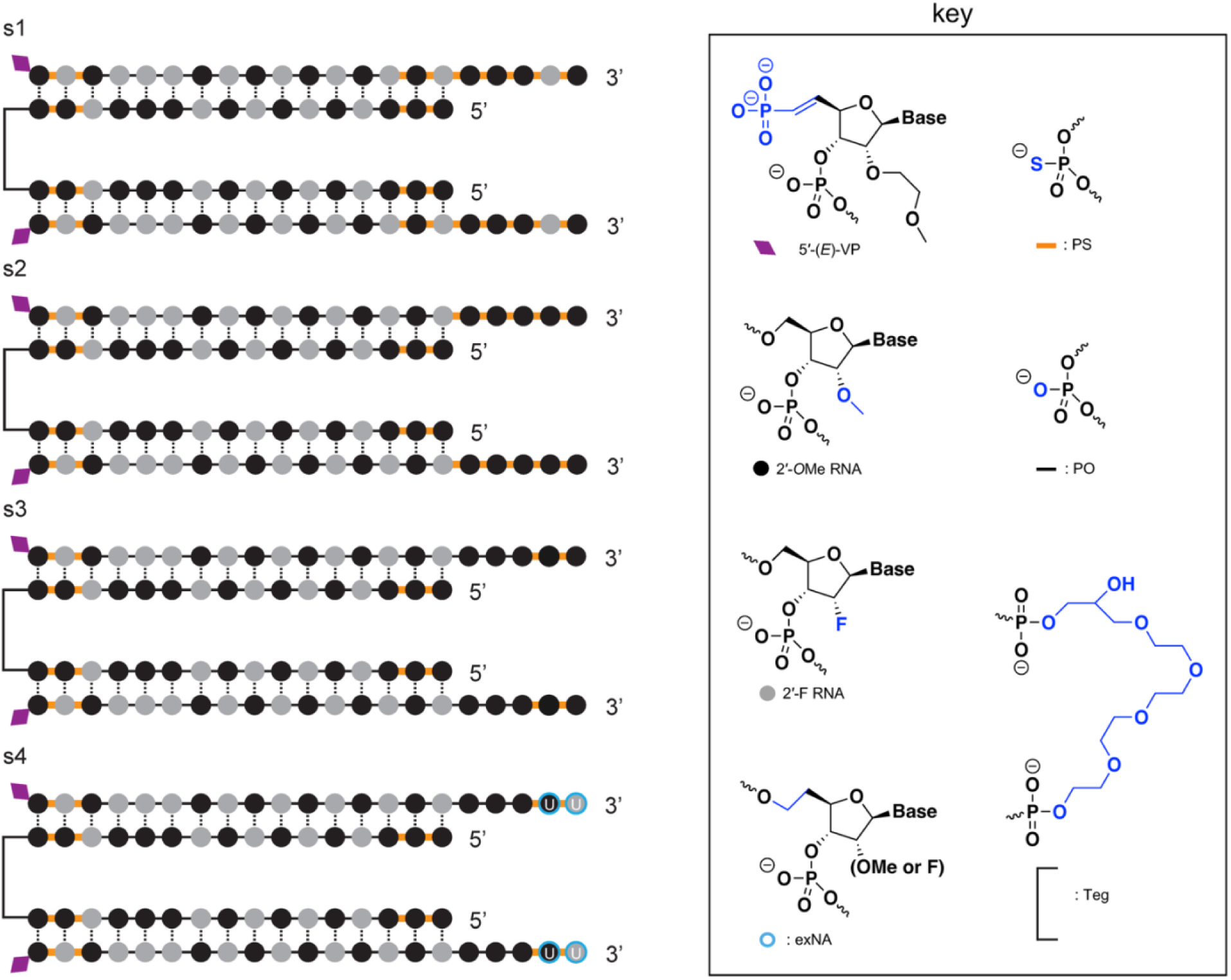
Chemical scaffolds used in this study. Abbreviations: 5′-(E)-VP, 5′-vinyl phosphonate; 2′-OMe, 2′-O-methyl; 2′-F, 2′-fluoro; exNA, extended nucleic acid; PS: phosphorothioate; PO: phosphodiester; TEG: tetraethylene glycol. The antisense strand of scaffold s4 was always synthesized with a fixed UU tail, hence the U displayed on the two 3′ nucleotides. Additional scaffolds that were used for screening and are only in supplementary figures are shown in Figure S1.

### Proof of concept in a prion disease model

We screened 20 siRNA sequences against mouse *Prnp* in mouse N2a cells, advancing 4 sequences into dose-response (55) (Figure S2). These studies nominated sequence 1682 (Table 1) as a tool compound for mouse *Prnp*. Sequence 1035, though inactive in that screen, was also tested in vivo based on its previously reported in cellulo activity in a different chemical scaffold (55) and its predicted cross-reactivity between mouse and human. Screening additional compounds in N2a cells by qPCR (Figure S3) yielded no additional strong hits. We tested 1682 and 1035 in wild-type C57BL/6N mice, not inoculated with prions, using a bilateral ICV bolus dose totaling 348 µg with tissue collection at 3-4 weeks post-dose and whole brain hemisphere total PrP quantified by ELISA (51) as a primary endpoint (Figure 2A). 1035 exhibited weak activity, with 81.8% residual PrP in the s1 scaffold (1035-s1), which improved with the s4 scaffold reaching 64.1% residual PrP. 1682-s4 was the most potent with 49.4% residual PrP, similar to previously reported ASO tool compounds (18, 51).

**Table 1.**
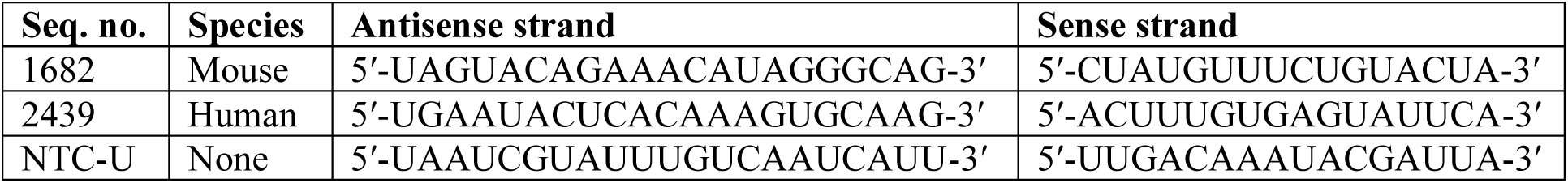
Properties of key siRNA sequences utilized. Note that where siRNAs were produced in the s4 scaffold, as indicated in Figure 1, the antisense strand always had a fixed UU 3ʹ tail regardless of the nucleotides shown in the sequence here. NTC-U denotes a non-targeting control sequence provided by UMass with no known target in the mouse or human genome. Sequences of all siRNAs tested in cellulo or in vivo are provided in the Supplementary Data.

**Figure 2.**
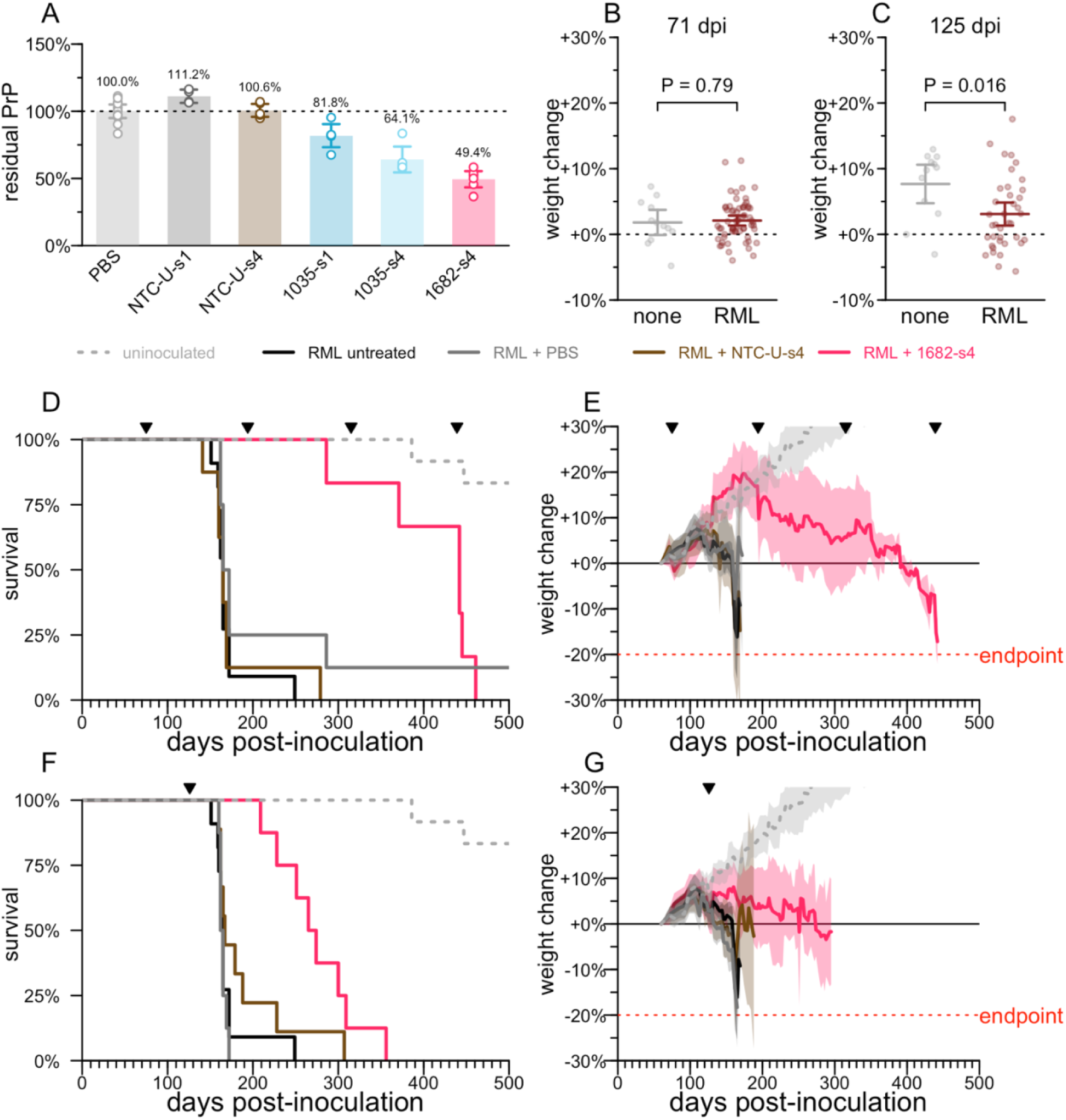
PrP lowering by divalent siRNA is effective in a mouse model of prion disease. **A)** Whole brain hemisphere PrP quantified by ELISA from 3-4 week target engagement studies in wild-type naïve mice receiving 348 µg of divalent siRNA, PBS or a non-targeting control (NTC-U). **B)** Weight change relative to individual animal baseline in animals at 71 days post inoculation (dpi), the last timepoint at which they were weighed prior to the 75 dpi intervention. At this timepoint, animals are asymptomatic and there is no difference in body weights (2-sided T test) between animals inoculated with Rocky Mountain Laboratories prions (RML) or uninoculated (none). Horizontal lines are are means and error bars are 95% CIs. **C)** Weight change relative to individual animal baseline in animals at 125 dpi, the last timepoint at which they were weighed prior to the 126 dpi intervention; this plot excludes animals treated beginning at 75 dpi. At this timepoint, animals are symptomatic, as evidenced by a significant difference in body weights (2-sided T test). Horizontal lines are are means and error bars are 95% CIs. **D)** Survival of animals in the early (75 dpi) intervention group. RML + PBS N = 7, RML + NTC-U-s4 N = 8, RML + 1682-s4 N = 6, RML untreated N=11, uninoculated N = 12. Ticks at the top show the dates of ICV drug administration. **E)** Weight trajectories of animals in the early (75 dpi) intervention group, expressed as percent change from each animal’s individual baseline. Solid lines are means, shaded areas are 95% confidence intervals. **F)** As in (D) but for the late (126 dpi) intervention group. RML + PBS N = 8, RML + NTC-U-s4 N = 9, RML + 1682-s4 N = 8; the RML untreated and uninoculated groups are repeated from panel D for reference. **G)** As in (E) but for the late (126 dpi) intervention group; the RML untreated and uninoculated groups are repeated from panel E for reference. For siRNA sequences see Table 1.

We intracerebrally inoculated wild-type mice with the RML strain of prions (50), which yields neuropathological changes detectable at the molecular level by ∼60 days post-inoculation (dpi) but no symptoms until at least 120 dpi (18). At 71 dpi, there was no difference in individual weight gain trajectory between inoculated mice and uninoculated controls, consistent with the presymptomatic disease stage (Figure 2B). In contrast, by 125 dpi, weight gain was significantly attenuated in the inoculated compared to uninoculated mice (P = 0.016), indicative of a symptomatic disease stage (Figure 2C).

Chronic dosing of 348 µg of 1682-s4 every 120 days (q120d) beginning at a presymptomatic timepoint of 75 dpi caused treated animals to significantly outlive controls by 2.7-fold (median 442 vs. 165 dpi, P = 0.0002, log-rank test, Figure 2D), with disease-attendant weight loss both delayed and slowed (Figure 2E). A single 348 µg dose given at a symptomatic timepoint of 126 dpi yielded survival time 3.5 months longer than controls (median 270 vs. 164 dpi, P = 0.0002, log-rank test, Figure 2F), with further weight loss delayed and slowed (Figure 2G). This difference amounts to a 64% increase in total survival time, or a 3.8x increase in remaining survival time from the moment of treatment at 126 dpi. All animals eventually succumbed to typical prion disease symptoms, and the majority met the pre-specified weight loss endpoint. Divalent siRNAs with a non-targeting control sequence (NTC-U), which did not lower PrP (Figure 2A), also did not increase survival time (Figure 2D, 2F), confirming on-target lowering of PrP as the mechanism of action.

These experiments confirmed that PrP lowering by divalent siRNA is effective against prion disease in mice.

### Generation of human *PRNP* transgenic mice

The observation of efficacy of PrP lowering by targeting the PrP RNA with divalent siRNA above led us to seek potent divalent siRNA compounds targeting the human *PRNP* gene. In vivo potency testing of genetically targeted human drug candidates, such as oligonucleotides, benefits from transgenic mice expressing the full human *PRNP* gene. Ours include 5’ and 3’ UTRs, coding sequence, and the sole intron.

We generated two new BAC transgenic lines harboring the full human *PRNP* gene, crossed them to homozygosity for endogenous *Prnp* knockout (ZH3/ZH3) (42), and determined their transgene integration sites, copy numbers, and human PrP expression level (Table 2, Figure 3). Tg25109 heterozygotes, with just 3 transgene copies, expressed human PrP at levels barely above wild-type (Figure 3A), while Tg26372 homozygotes, with 20 transgene copies, expressed human PrP at 5.4-fold the wild-type level (Figure 3A). The relationship between transgene DNA copy number and protein expression was sub-linear, just as observed (13) for transgenes encoding mouse PrP (Figure 3B). Short-read sequencing of the Tg26372 mouse confirmed integration of 46.0 kb of human sequence from the major and ancestral 129M haplotype: 20.5 kb upstream, 15.2 kb spanning from the *PRNP* transcription start to stop site, and 10.3 kb of downstream sequence (Figure 3C). The transgene excludes the downstream gene *PRND*, for which overexpression in the brain is known to be toxic.

**Table 2.** Transgenic human PRNP mice. Each line harbors a tandem array of the same 46.0 kb bacterial artificial chromosome (BAC) containing human PRNP 129M, randomly integrated into a different site in the mouse genome, see Methods for details. All mice are on a C57BL/6N background and PrP expression level is reported as a fold change relative to WT C57BL/6N mice, n = 3-6 per group. *Tg25109 homozygotes were subviable, see Results for details. The in-house ELISA assay used for these protein expression analyses has been shown equally reactive for mouse PrP and human PrP (51).

| Line | Integration site | Genes disrupted at integration site | Het copy number | Het PrP expression (fold WT) | Hom copy number | Hom PrP expression (fold WT) |
| --- | --- | --- | --- | --- | --- | --- |
| Tg25109 | chr12 | *Frmd6, Tmx1* | 3 | 1.1 | 6* | 2.0* |
| Tg26372 | chr18 | *Dok6* | 10 | 3.4 | 20 | 5.4 |

**Figure 3.**
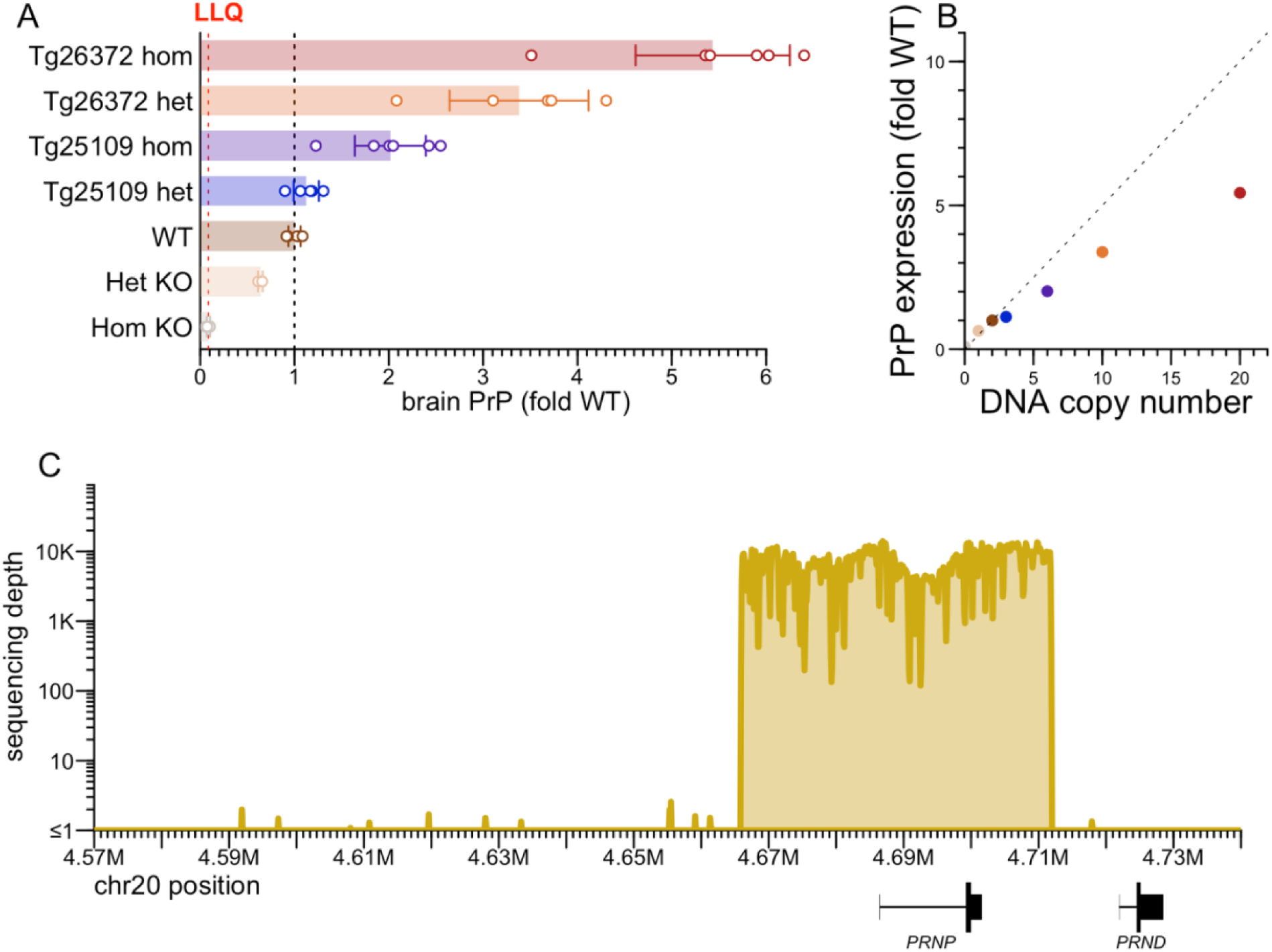
Characterization of human PRNP BAC transgenic mice. **A)** PrP concentration in whole brain hemispheres, normalized to the mean of wild-type (WT) animals. Het and hom KO animals are of the ZH3 PrP knockout line (42). Points are individual animals. N = 3-6 per group. Rectangular bars are means. Error bars are 95% confidence intervals. The lower limit of quantitation (LLQ) is 10 ng/g based on a 0.05 ng/mL bottom standard curve point and a 1:200 dilution of brain homogenate. **B)** PrP expression from (A) versus gene copy number at the DNA level; the dashed line with slope 0.5 represents the linear relationship whereby 2 gene copies = 1-fold expression. **C)** Extent of human sequence in the bacterial artificial chromosome (BAC), based on targeted capture sequencing (Methods) with reads aligned to the human genome reference (GRCh38), using genomic DNA from a Tg26372 mouse.

Heterozygote-heterozygote crosses of our Tg25109 line yielded a ratio of 98:198:32 (non-transgenic:heterozygote:homozygote), a significant deviation from Mendelian ratio (P < 1e-15, Chi-square test). Of 4 Tg25109 homozygous x homozygous pairs mated, only 1 pup was ever born. We examined the literature evidence for viability of knockouts for *Tmx1* and *Frmd6*, the two genes disrupted at the Tg25109 integration site (Table 2). *Tmx1* (formerly known as *Txndc1*) homozygous knockout mice were reported to have abnormal bone metabolism and immunology (56), while *Frmd6* homozygous knockout mice were reported to have several phenotypes including eye and hematological defects and were born at less than the expected Mendelian ratio from het-het crosses (57, 58) (41:51:15, P = 0.0016, Chi-square test). The sub-viability of Tg25109 homozygous mice therefore likely arises from knockout of *Frmd6,* or from the combination of both *Frmd6* and *Tmx1*. It is unlikely to be transgene expression-related, as Tg26372 homozygous mice, with higher copy number (20 vs. 6) and protein expression (5.4-vs. 2.0-fold wild-type) were unaffected. Given the difficulty of obtaining adequate numbers of Tg25109 homozygotes for experiments, we excluded this genotype from further experiments. Given the convenience of maintaining Tg26372 as obligate homozygotes, we performed the vast majority of experiments in this genotype, but 2 compounds also tested in Tg25109 heterozygotes showed target engagement in both genotypes (Figure S4).

These mice provided us a model in which to develop siRNA compounds against the human *PRNP* RNA.

### Identification of compounds targeting human *PRNP*

A screen of 24 predicted active compounds in human A549 cells identified 4 hits with dose-responsive potency, led by sequence 2440 (Figure S2) (55). We conducted an expanded screen of 84 compounds in human U251-MG glioblastoma cells, prioritizing predicted hot spots within a ±3 base pair walk of the top sequences from the initial screen (Figure 4A). At a 2.0 µM screening concentration, 34 compounds (40%) yielded <10% residual *PRNP*. The 0.5 µM screening concentration provided better power to discriminate the most potent sequences, with just 14 (17%) yielding <10% residual *PRNP* (Figure 4A). 10 sequences tested in dose-response all proved active with IC_50_ < 100 nM (Figure 4B).

**Figure 4.**
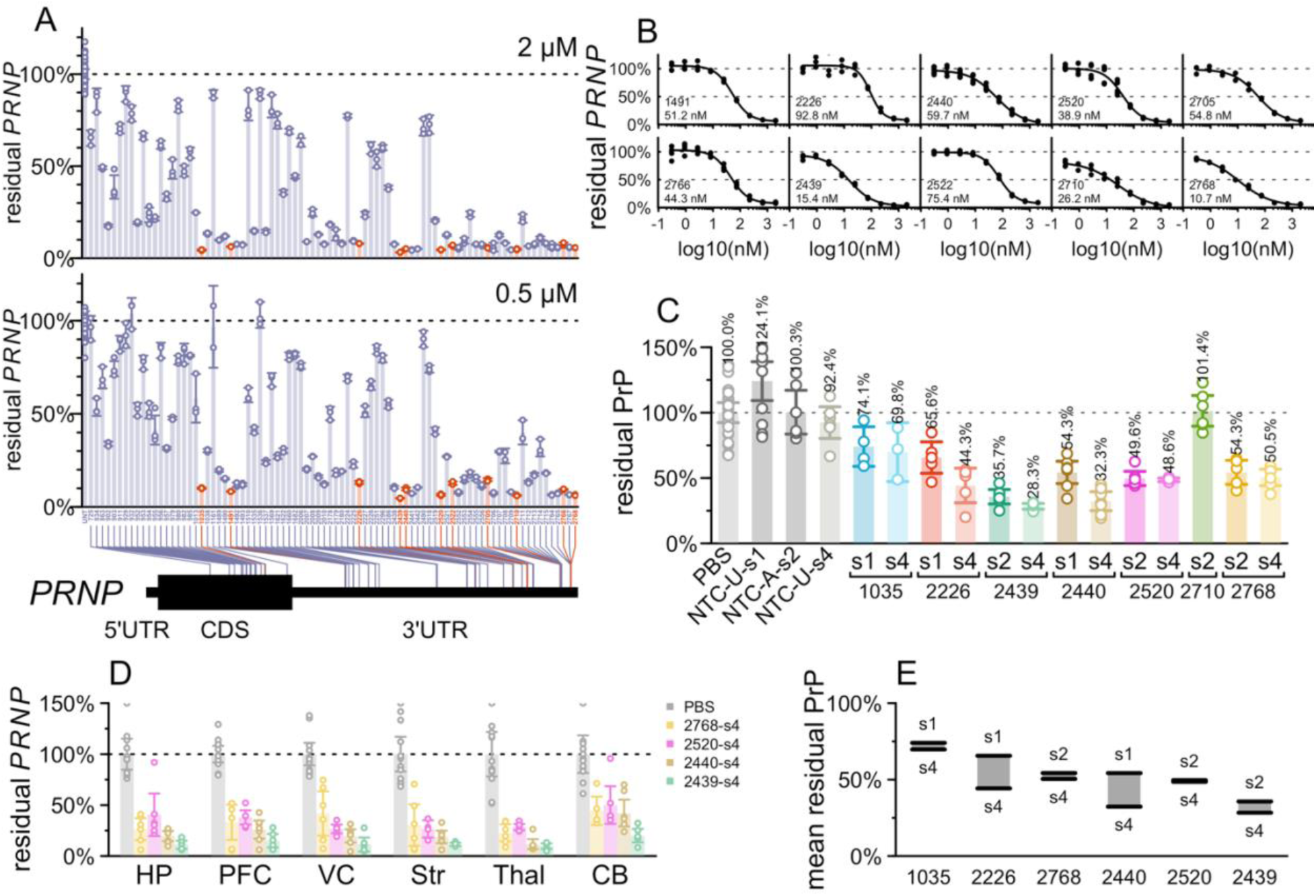
Identification of human PRNP-targeting siRNA sequences. **A)** In cellulo screening using scaffold Chol-TEG s2 (Figure S1). The readout is PRNP qPCR, normalized to the mean of untreated (UNT) controls. Each point represents 1 well of U251-MG human glioblastoma cells, run in qPCR technical duplicate. Rectangular bars represent the mean of triplicate wells, and error bars represent the 95% confidence intervals. Screening used Chol-TEG conjugated monovalent siRNAs (Figure S1) by gymnotic uptake at 2.0 µM (top panel) or 0.5 µM (bottom). N = 3 per group. **B)** Dose-response testing in cell culture. The readout is PRNP qPCR. Points represent individual wells of U251-MG cells, curves are four-parameter dose-response curves fit by the drc package in R. Displayed at bottom are the sequence number and the calculated IC_50_ value. N = 3 per group. **C)** In vivo testing. The readout is PrP ELISA. Each point represents a whole brain hemisphere of one Tg26372 homozygous mouse, rectangular bars represent the group mean, and error bars represent the 95% confidence intervals. N = 3-7 per group, total 120 animals. **D)** Regional PRNP qPCR for select compounds from the same animals in panel (C), in mouse hippocampus (HP), prefrontal cortex (PFC), visual cortex (VC), striatum (Str), thalamus (Thal), and cerebellum (CB). Rectangular bars represent the group mean, and error bars represent the 95% confidence intervals. N=5-6 per group. **E)** Data replotted from panel (C) — difference between mean residual PrP for the s1/s2 (high PS, no exNA) scaffolds versus the s4 (reduced PS, with exNA) scaffold for 6 sequences where both were tested in vivo. For siRNA sequences see Table 1 and Supplementary Data.

7 sequences selected based on potency as well as cross-reactivity were advanced to in vivo screening in multiple chemical scaffolds (Figure 1, Figure 4C) alongside PBS and non-targeting controls (NTCs), for a total of 17 experimental groups totaling 120 Tg26372 homozygous animals. Each compound was tested at a 348 µg dose, with whole brain hemispheres collected at 4-5 weeks post-dose analyzed by ELISA. Non-targeting controls did not significantly lower PrP in any chemical configuration tested. In either chemical scaffold, 2439 (Table 1) proved to be the most potent sequence in vivo (Figure 4C). RT-qPCR analysis for the top 4 sequences tested in the s4 scaffold confirmed that 2439 achieved deeper *PRNP* RNA lowering than the other 3 sequences in 6/6 mouse brain regions tested (Figure 4D). For all compounds, lowering was weakest in the cerebellum, as reported for a divalent siRNA targeting *HTT* (28), likely due to lower drug accumulation in rodent cerebellum. The deeper lowering at the RNA level (residual ranging from 9.6% in thalamus to 19.9% in cerebellum) than at the protein level (28.3% residual in whole hemisphere) in this experiment may simply reflect floor effects in our ELISA: a contrived sample designed to mimic 10% residual PrP (10% wild-type brain homogenate mixed with 90% knockout brain homogenate) read out as an average of 17% of wild-type across all ELISA plates in this study (Figure S6). For 6/6 sequences where both a high-phosphorothioate (s1 or s2) and the low-phosphorothioate plus exNA-containing s4 scaffold were tested, s4 was numerically the more potent, by a margin of 1 to 21 percentage points of PrP lowering (Figure 4E), as shown for *Htt* and *Apoe* (34). This difference was observed for s4 versus both s1 and s2, though a qualifier is that due to differences in accounting for hypochromicity of duplexed siRNA, the administered dose of s2 may have been 14% lower (see Methods).

We used tribromoethanol as an anesthetic for our initial studies because it was used for divalent siRNA previously (28). We later pivoted to 3% isoflurane anesthesia with incorporation of divalent cations (a 14:2:1 molar ratio of Ca^2+^:Mg^2+^:divalent siRNA) into the injectable solution, which has been reported to eliminate seizures upon injection of high-dose oligonucleotides into CSF (45). All animals recovered from anesthesia normally, we never observed seizures. Animals generally gained weight for the duration of the in-life period (Figure S5), with the exception of compound 2520-s2, for which 5/6 animals experienced acute weight loss between 3 and 4 weeks post-dose. To further assess tolerability, we performed RT-qPCR for neuroinflammatory markers *Gfap* and *Iba1* in the visual cortex. None of the sequences tested significantly affected *Gfap* (all *P* > 0.10, Dunnett’s test); only 2520-s4 marginally impacted *Iba1* (43% decrease, P = 0.046, Dunnett’s test; Figure S5). 2439 exhibited a favorable in silico predicted off-target profile: antisense strand bases 2-17 harbored at least 2 mismatches to all human protein-coding mRNAs other than *PRNP* (Supplementary Data).

These studies support the selection of 2439 as the lead sequence.

### Comparison of chemical scaffolds and antisense strand 3′ tails

The in cellulo and in vivo screening results supported the selection of 2439 as the lead sequence (Figure 4C) and provided some evidence that the s4 scaffold resulted in a deeper knockdown than the s1 and s2 scaffolds (Figure 4E), but we sought additional evidence to confirm the lead scaffold before proceeding. Our s4 compounds were all synthesized with a fixed 3′-UU tail mismatched to the target RNA, initially due to the relative ease of synthesis of 2′OMe-exNA-uracil (mxU) and 2′F-exNA-uracil (fxU) phosphoramidites (34) and the unavailability of their A, G, or C equivalents. In contrast, our s1 and s2 compounds were synthesized with full complementarity to the target RNA. Depending upon sequence, unmatched 3′ tails can facilitate PAZ binding (59) without compromising RISC activity (60, 61). Complete complementarity has even been associated with increased rates of RISC unloading (35) and target-directed degradation (36). We therefore sought to determine for the lead sequence the relative contributions of the PS and exNA modifications that distinguish the s4 scaffold, versus the effect of this 3′ fixed tail.

We performed a 3-point in vivo dose response experiment with 5-fold dose increments (7, 35, and 174 µg) for each of 3 scaffolds (s2 matched tail, s2 fixed tail, s4 fixed tail) versus PBS controls, N = 8 per group, with harvest at 4 weeks. We also included s3 fixed tail at only the highest dose (174 µg) to test our assumption that reduction of PS content without introduction of exNA would result in reduced activity.

At every dose level, 2439-s4 was the superior compound in terms of whole-hemisphere PrP, with 29.5% residual at the 174 µg dose (Figure 5A). As expected, the s3 scaffold with fixed tail had less knockdown than any other scaffold at the high dose (Figure 5B). We performed RT-qPCR for *PRNP* RNA and fit dose-response curves for 6 brain regions (Figure 5C), using a linear model (Methods) to characterize the effects of scaffold/tail combination, brain region, and dose level. This model indicated that s1 fixed tail provided 9.4 percentage points deeper *PRNP* lowering than s1 matched tail (P = 0.0017), while the s4 fixed tail conferred another 15.9 percentage points beyond s1 fixed tail (P = 5.0e-11, Figure 5D). Thus, both the fixed tail and the terminal exNA modification contributed to the potency of the s4 scaffold. At the middle dose (35 µg), 2439-s4 yielded <50% residual *PRNP* in 6/6 brain regions tested (Figure 5C), with target engagement weakest in cerebellum (Figure 5E) as expected (28). Individual dose-response curves for each scaffold/tail combination in each brain region yielded median effective dose (ED_50_) for 2439-s4 ranging from 5 µg in thalamus to 18 µg in cerebellum.

**Figure 5.**
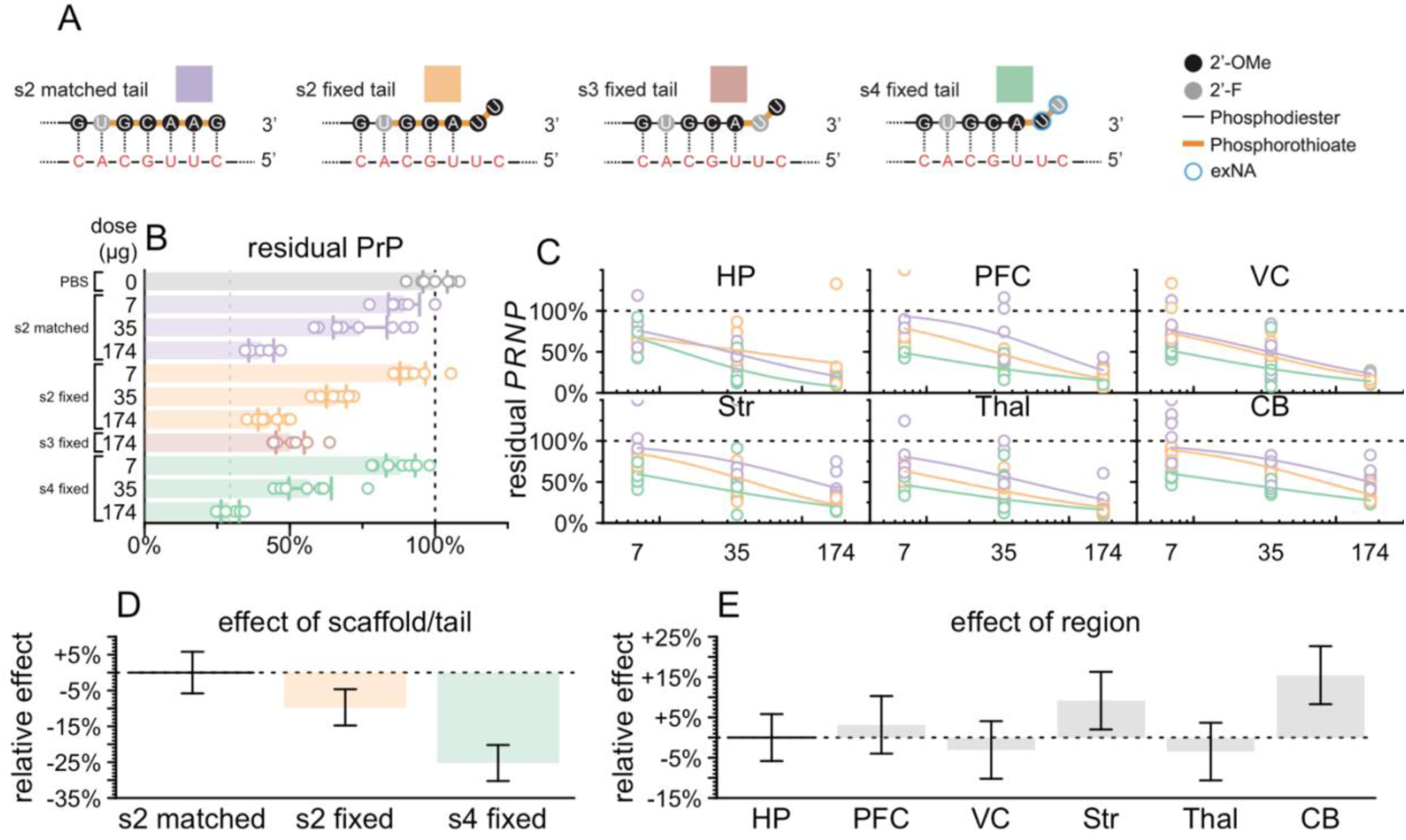
Impact of chemical scaffold and 3′ antisense tail on lead sequence potency. **A)** Color key and diagram of differences between 4 compounds tested. These compounds differ only in the 3′ tail of the antisense strand shown here. The antisense strand of the divalent siRNA is shown at top, and the target mRNA is shown in red below. Each compound was injected at the doses indicated in B-C into N = 8 animals harvested after 30 days. **B)** Whole hemisphere residual PrP by ELISA (x axis) by compound and dose (y axis). Each point is one animal, rectangular bars are means, error bars are 95% confidence intervals. **C)** Regional PRNP RNA by qPCR (y axis) versus dose (x axis). Each point is one animal. Curves are 4-parameter log-logistic dose-response curves (see Methods). Brain regions analyzed: hippocampus (HP), prefrontal cortex (PFC), visual cortex (VC), striatum (Str), thalamus (Thal), and cerebellum (CB). **D)** Linear model coefficients for scaffolds fit to the data in (C). Rectangular bars are means, error bars are 95% confidence intervals. **E)** Linear model coefficients for brain regions fit to the data in (C). Rectangular bars are means, error bars are 95% confidence intervals.

This experiment confirmed 2439-s4 with its fixed 3′-UU tail (full structure in Figure S7) as our drug candidate.

### Durability and dosing regimens for drug candidate

We tested the potency and durability of effect of 2439-s4 (Figure 6A) in additional studies. Pharmacodynamic effect measured by RT-qPCR of *PRNP* mRNA in whole brain hemisphere at 30 days post-dose showed dose-responsive target engagement over a two order of magnitude range both in administered dose and in drug accumulation in tissue (Figure 6B, Figure S8A-B). Fitting a PD/PK model returned a tissue IC_50_ value of 1.2 µg/g, with generally 1-2% of administered dose retained in brain 30 days post-dose (Figure S8C). Single 348 or 174 µg doses yielded activity out to at least 4 months (Figure 6C) and 6 months post-dose (Figure 6D), by measuring PrP protein by ELISA. The same sequence in 2 scaffolds with higher PS content provided superior durability but lower initial knockdown at the 1-month timepoint (Figure S9).

**Figure 6.**
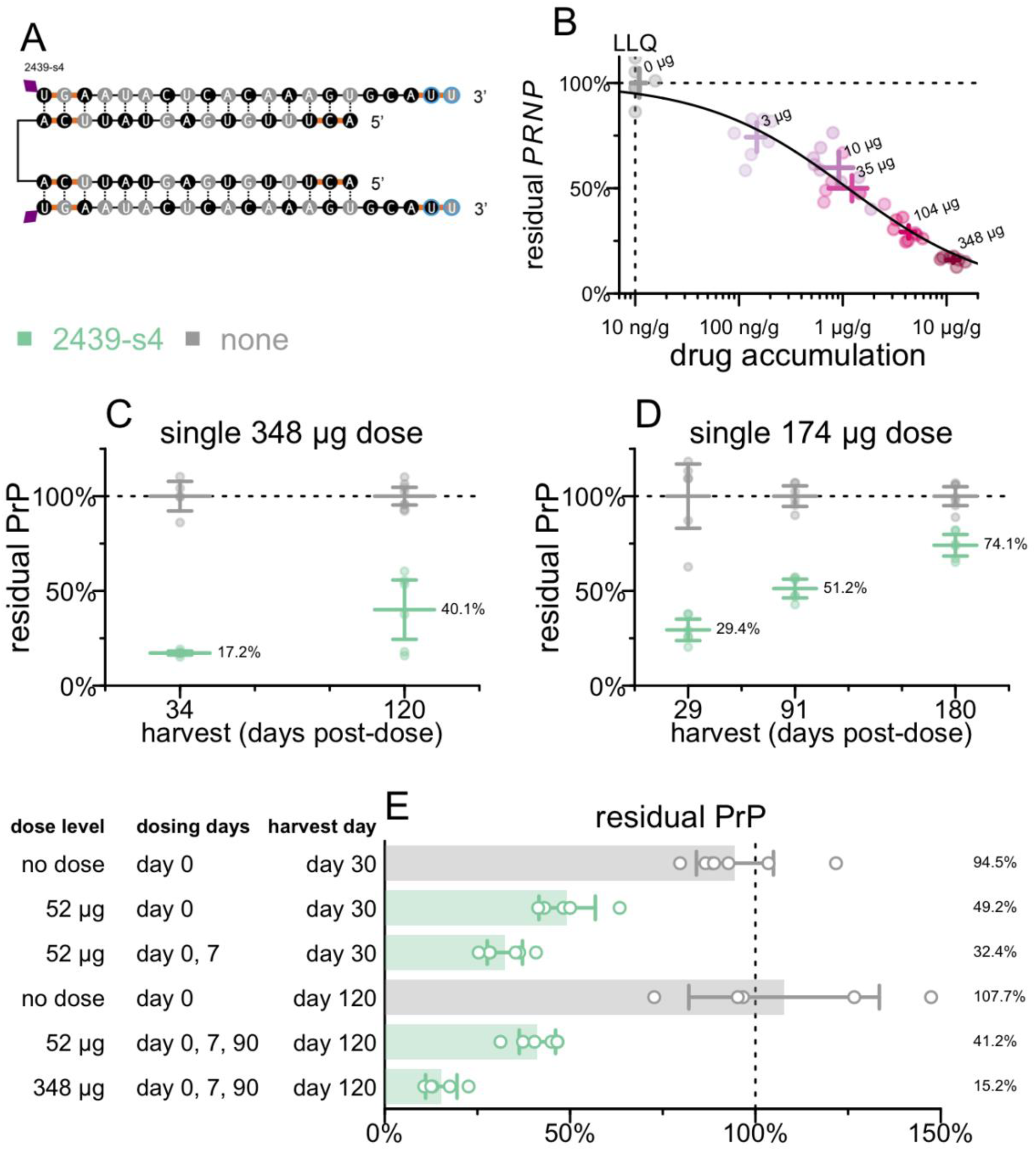
Characterization of lead compound 2439-s4. **A)** Identity of 2439-s4. **B)** Dose-responsive effect of 2439-s4 on PRNP mRNA in whole hemisphere. N = 6-8/group. **C)** Durability of effect of a single 348 µg dose in Tg26372 animals. Whole hemisphere PrP (y axis) versus months post-dose (x axis). Each point is one animal (N = 4-8/group), horizontal line segments are means, error bars are 95% confidence intervals. Harvest days are exact. **D)** Durability of effect of a single 174 µg dose in Tg26372 animals. Whole hemisphere PrP (y axis) versus months post-dose (x axis). Each point is one animal (N = 6/group), horizontal line segments are means, error bars are 95% confidence intervals. Harvest days are exact. **E)** Impact of repeat dosing regimens on target engagement in Tg26372 animals. Whole hemisphere PrP (x axis) for indicated dose levels and regimens (y axis). Each point is one animal (N = 5-6/group), rectangular bars are means, error bars are 95% confidence intervals. Harvest days indicated are ±3 days.

We also investigated a loading dose strategy, with a second dose given 7 days after the initial dose, and of repeat dosing after 90 days (Figure 6E). We evaluated both the original 348 µg dose as well as a 52 µg dose which corresponds to a dose reachable clinically (see Discussion). Compared to a single dose at day 0, a loading dose regimen (day 0 and 7) for 52 µg provided improved target engagement at day 30, with 5/6 regions below 25% residual *PRNP* RNA (Figure S10A); all regions were below the respective levels reached with the 348 µg dose of 1682-s2 (Figure S10B) in the survival study (Figure 2).

These experiments indicated that a single dose of divalent siRNA can provide target engagement and durability expected to be meaningful on the time course of prion disease, while repeat dosing further improves target engagement.

### Investigational New Drug (IND)-enabling studies

We produced a batch of 2439-s4 under Good Manufacturing Practices (GMP) for IND-enabling toxicology and future clinical studies (Figure S11). In single-dose intrathecal toxicology studies, dogs receiving 0, 20, 60, or 200 mg and rats receiving 0, 0.3, 1, or 3 mg 2439-s4 were necropsied at 1 day post-injection and 28 days post-injection; no adverse findings were discovered at any dose in either species (Table 3). Plasma exposure peaked at 0.5 – 4 hours post dose; drug concentrations measured in CSF were highly variable and not predictive of brain parenchyma exposure (Figure S12). Drug exposure in brain was well above the estimated IC_50_ of ∼1.2 µg/g (Figure 6B) across spinal cord, cerebellum, and cortex in both species (Figure 7A-B), whereas it was below this IC_50_ in the deepest brain regions, particularly in dog (Figure 7A). Approximately 1% of administered dose was retained in the brain (Table 4). There were no effects on hematology, coagulation, or serum chemistry attributed to administration of 2439-s4 in rats or in dogs.

**Table 3.** Summary of IND-enabling GLP toxicology study results. RoA: route of administration. IT: intrathecal. M: male, F: female. Findings: moribundity and mortality, cage-side clinical observations, weekly body weights, food consumption, ophthalmic exams, noninvasive homecage neurobehavioral assessment (dogs only), electrocardiogram (ECG)(dogs only), noninvasive blood pressure (dogs only), respiratory rate (dogs only), hematology, serum chemistry, and coagulation, urinalysis, and histopathology.

| Species | RoA | Doses | In-life period | Dose level (mg) | N | Findings |
| --- | --- | --- | --- | --- | --- | --- |
| Rat | IT | 1 | 1 day (core) | 0 | 10M/10F | None |
|  |  |  |  | 0.3 | 10M/10F | None |
|  |  |  |  | 1.0 | 10M/10F | None |
|  |  |  |  | 3.0 | 10M/10F | None |
| Rat | IT | 1 | 28 day (recovery) | 0 | 5M/5F | None |
|  |  |  |  | 0.3 | 5M/5F | None |
|  |  |  |  | 1.0 | 5M/5F | None |
|  |  |  |  | 3.0 | 5M/5F | None |
| Dog | IT | 1 | 1 day (core) | 0 | 4M/4F | None |
|  |  |  |  | 20 | 4M/4F | None |
|  |  |  |  | 60 | 4M/4F | Non-adverse gliosis, 1/8 |
|  |  |  |  | 200 | 4M/4F | Non-adverse gliosis, 3/8 |
| Dog | IT | 1 | 28 day (recovery) | 0 | 2M/2F | None |
|  |  |  |  | 20 | 2M/2F | None |
|  |  |  |  | 60 | 2M/2F | None |
|  |  |  |  | 200 | 2M/2F | None |

**Figure 7.**
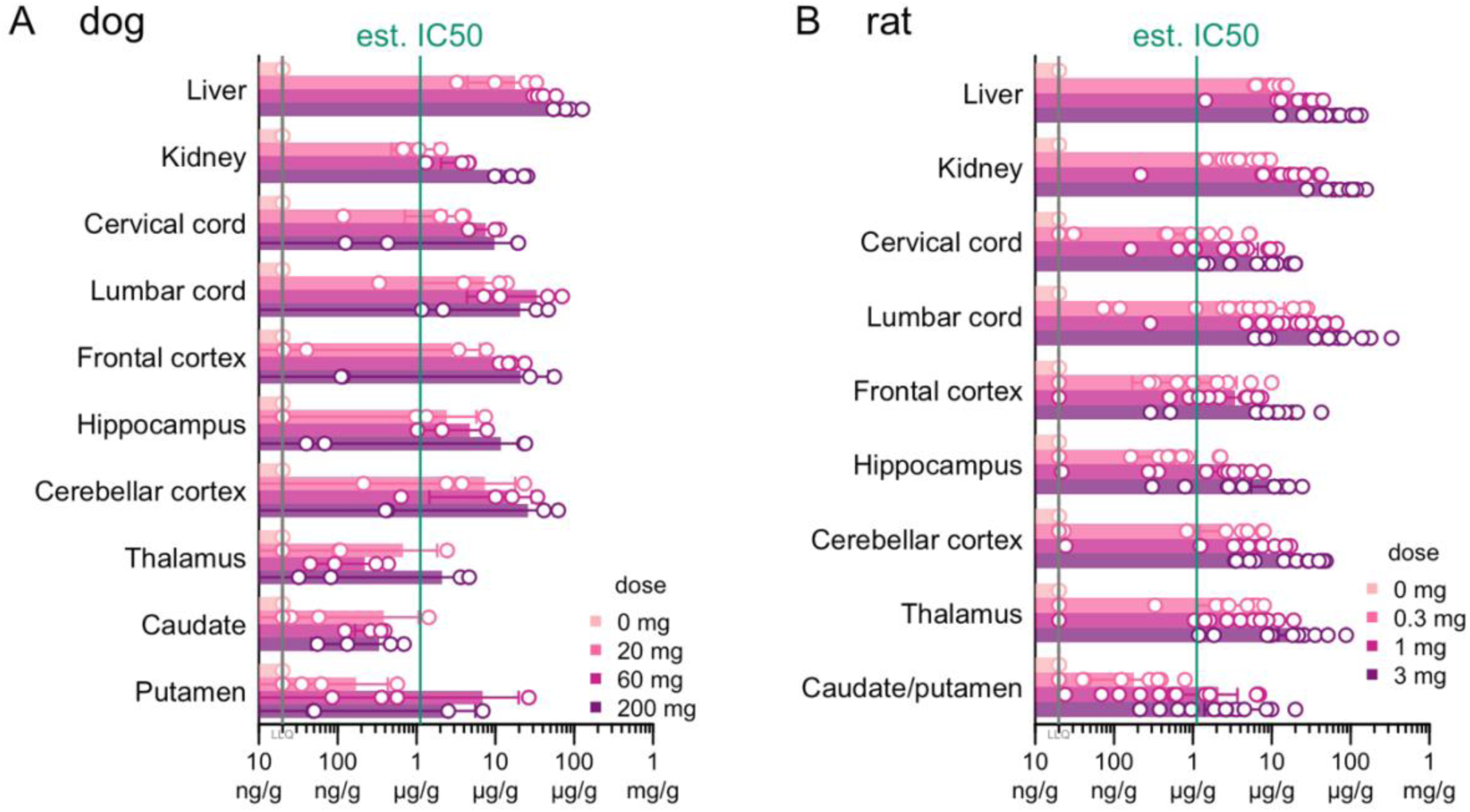
Biodistribution of intrathecally delivered 2439-s4 in Good Laboratory Practices toxicology studies. Tissue concentration of 2439-s4 in **A)** dogs (N = 2 male/2 female per group) 28 days after and **B)** rats (N=6 male/6 female per non-zero group, 3 male/3 female for zero dose) 3 days after a single intrathecal dose of 2439-s4 at the indicated doses. Day 3 tissue was collected from toxicokinetic (TK) cohort rats. The estimated IC_50_ of 1.2 µg/g was obtained from the pharmacodynamic/pharmacokinetic (PD/PK) model in Figure 6B

**Table 4.** Proportion of administered dose retained by tissue, species, and days post-dose. Proportions are averaged across 3 dose levels (low, medium, and high doses as shown in Figure 7), with 4 male (M)/4 female (F) dogs per dose at day 1, 2M/2F dogs at day 28, and 6M/6F rats at day 3. Day 3 measurements are from the toxicokinetic (TK) cohort blood draws.

| species<br>days post-dose | dog |  | rat |
| --- | --- | --- | --- |
|  | 1 | 28 | 3 |
| **Brain** | 1.1% | 1.0% | 1.6% |
| **Kidney** | 0.9% | 0.3% | 4.3% |
| **Liver** | 42.0% | 14.5% | 23.2% |

Cyp inhibition studies using human hepatocytes found no interactions with IC_50_ <400 nM. Cyp induction studies using human hepatocytes found no cytotoxic effect of 2439-s4 on cells and no effect on Cyp enzyme expression levels at concentrations up to 400 nM. Transporter inhibition studies in Caco-2 cells (P-gp), MDCKII cells (BCRP), or HEK293 cells found a maximum of 27.2% inhibition of OAT3 in the presence of 2439-s4 concentrations up to 400nM. 2439-s4 was not an inhibitor of any other transporters tested under the conditions tested. Transporter substrate studies in MDCKII cells found that 2439-s4 was not a substrate of P-gp or BCRP under the conditions tested.

No evidence of genotoxicity was found using the in vitro micronucleus test in CHO-K1 cells at concentrations up to 500 μg/mL in the presence or absence of metabolic activation. No evidence of mutagenicity was found using the Ames test in five *Salmonella* strains or one *E. coli* strain at concentrations up to 5000 μg/plate in the presence or absence of metabolic activation.

The above results supported advancing 2439-s4 to clinical studies, and an Investigational New Drug application based on these data was cleared by the U.S. Food and Drug Administration.

## Discussion

Our study shows that lowering PrP RNA with a divalent siRNA is effective against prion disease, and that the human drug candidate, 2439-s4, is potent, long-lasting, and appears well-tolerated in single dose nonclinical toxicology studies.

As with ASOs (17, 18), we observed an extension of healthy life in prion-infected animals treated pre-symptomatically with divalent siRNA, whereas in already-symptomatic animals treatment extended life without reversing existing weight loss. This is consistent with an inability of PrP lowering to address pre-existing neuronal loss, and with a lag time of a few weeks between target engagement at the RNA level and maximal lowering of PrP at the protein level (49). Our data support treatment of prion disease patients at both symptomatic and pre-symptomatic timepoints, while suggesting that the greatest good can be achieved in a pre-symptomatic preventive paradigm (2).

The drug candidate 2439-s4 appears to have favorable properties in terms of potency and durability. We observed a depth of target suppression — as low as 17% residual whole hemisphere PrP after a single 348 µg dose and 15% with a loading dose followed by repeat dosing — never previously reported for PrP, and we showed at least some activity out to 6 months after a single dose. The ED_50_ for the candidate in mice, measured by regional qPCR, ranges from 5 - 18 µg depending upon brain region, which compares favorably to the ED_50_ values ranging from 27 µg (in spinal cord) to 69 µg (in cortex) reported for the most potent human ASO candidate against PrP (62). Using CSF volume scaling (0.04 mL in mice versus 130 mL in human (63, 64), a factor of 3,250),18 µg might correspond to 58 mg in a human. This dose level is clinically precedented for oligonucleotides — the ASO tofersen for *SOD1* ALS is dosed at 100 mg (65). These calculations lead us to hypothesize that a single dose of 2439-s4 could lower PrP by 50% in many human brain regions, which is important because prion disease is a whole brain disease. A limitation of our study, however, is that uniformity of brain exposure is a major challenge for any intrathecally delivered oligonucleotide therapy (66, 67). 2439-s4 is not sequence-matched to rat or dog *PRNP*; we did not assess biodistribution and target engagement in a pharmacodynamically relevant toxicology species. Divalent siRNAs dosed into non-human primates or sheep by ICV or IT routes (28, 68, 69) were reported to have broad distribution and activity, although, as with ASOs (66), deep subcortical brain regions are less well-reached, with drug concentration in putamen being <20% that achieved in cortex.

A limitation of our study is that our comparison of siRNA chemical scaffolds focused on one sequence and may not be generalizable. Moreover, due to the hypochromicity of duplexed siRNAs, the administered doses of different scaffolds may not be identical, urging caution around interpretation of these scaffolds’ relative potency. Another limitation of our study is that, although we demonstrated efficacy in a disease model using a tool compound, we did not assess the disease modifying impact of the deeper PrP lowering achieved with our clinical candidate. Recent reports of researchers dying of prion disease after occupational exposure to human prion-infected brain tissue (70, 71) convinced us to not examine the efficacy of our drug candidate in a challenge study with human prions in our humanized mice. Our drug candidate is not cross-reactive for mouse *Prnp*, precluding rescue studies in wild-type mice. Thus, we were unable to measure the survival benefit attainable by the deeper PrP lowering observed with our drug candidate compared to our mouse *Prnp* tool compound. Delay of prion disease by PrP lowering is dose-responsive (18), and homozygous knockouts are invulnerable to prions (11), anchoring our assumption that deeper lowering is better. Nevertheless, at present we lack an animal model system to answer in a more quantitative way what benefit will be achieved by the deep PrP lowering described here — for instance, whether prion replication or symptom progression could be halted.

GLP toxicology studies did not find any adverse effects attributable to 2439-s4 in rats or dogs. Drug-drug interaction studies did not find evidence of cytochrome P450 inhibition with an IC_50_ < 400 nM or cytochrome P450 induction at concentrations up to 400 nM. Transporter inhibition studies found no evidence of inhibition with an IC_50_ < 400 nM. Genotoxicity studies found no evidence of genotoxicity or mutagenicity under the conditions tested.

Based on the studies described here, we have filed an IND with the U.S. Food and Drug Administration and received clearance to initiate a single-dose clinical trial in prion disease patients (ClinicalTrials.gov NCT07444580). Given the severity and rapid progression of prion disease, we asked FDA for permission to proceed to clinical studies based on rodent toxicology alone but were advised that two species would be a requirement, thus leading to the rat and dog toxicology performed here. Our interactions with FDA identified several ways to reduce the number of drug product vials required for quality control and stability testing, thus making production of a small GMP batch feasible for clinical studies. Because FDA evaluates every program individually, our regulatory interactions may not be predictive of what will be permitted for other novel modalities in similarly rare and severe diseases. Nevertheless, we have made the text of our IND and our interactions with FDA publicly available (see Data Availability statement) as a service to the rare disease community.

Other potential PrP-lowering drugs reported include an intrathecal ASO (18), intravenous one-time viral vectored base editors (19) and zinc finger suppressors (72), and chronically orally administered small-molecule molecular gates blocking PrP’s transit through the Sec61 translocon (73).Of these candidates, only an ASO has reached clinical trials. Divalent siRNA 2439-s4 offers another potential therapeutic ready for clinical application.

## Data Availability

Raw individual-level animal data, source code sufficient to reproduce all analyses herein, and the text our Investigational New Drug application and regulatory interactions with U.S. FDA are available in this study’s online git repository: https://github.com/ericminikel/divalent

## Supplementary Data Statement

Supplementary Data are available at NAR Online.

## Acknowledgments

We thank the following scientists who contributed to IND-enabling studies: Danielle Breslow, Dave Butler, Rebecca Campbell, Josh De Los Santos, Chris Duffy, Steve Folio, Nichole Franz, Seth Gibbs, Beibei Guo, Richard Hargis, Melissa Harned, Lois Haupt, Jennifer Horkman, Thessa Jacob, Niels King, Lily Li, Brian Lu, Rambabu Naravaneni, Sonja Neitzel, Eric (Yingchao) Niu, Varsha Paradkar, Frank (Wanping) Rao, Nicole Rottman, Sarah Shellenbarger, Jeremy Smith, Wangqiyue Sun, Yansheng Wu, Yuwei Xiao, Jim (Jimin) Yang, Wenhao Yao, Xinyu Zhang.

## Author Contributions Statement

Conceptualization: JEG, ALJ, JFA, EVM, AK, SMV

Data curation: JEG, EVM

Formal analysis: JEG, EVM

Funding acquisition: EVM, AK, SMV

Investigation: JEG, TLC, FES, DE, ZEK, CLG, MRH, GAK, MNK, NGK, YL, KYG, RM, KD, MH, KG, NC, VL, DTC, ALJ, KY, JFA, EVM, AK, SMV,

Methodology: JEG, TLC, FES, DE, ZEK, MRH, GAK, KF, ML, AAC, EVM, AK, SMV

Project administration: ALJ, AAC, EVM, AK, SMV

Resources: DE, ZEK, CLG, MRH, GAK, KF, ML, ALJ, KY

Supervision: EVM, AK, SMV

Visualization: JEG, FES, MNK, YL, EVM

Writing – original draft: EVM

Writing – review & editing: all authors

## References

1. Prusiner, S.B. (1998) Prions. Proc. Natl. Acad. Sci. U. S. A., 95, 13363–13383.

2. Vallabh, S.M., Minikel, E.V., Schreiber, S.L. and Lander, E.S. (2020) Towards a treatment for genetic prion disease: trials and biomarkers. Lancet Neurol., 19, 361–368.

3. Büeler, H., Fischer, M., Lang, Y., Bluethmann, H., Lipp, H.P., DeArmond, S.J., Prusiner, S.B., Aguet, M. and Weissmann, C. (1992) Normal development and behaviour of mice lacking the neuronal cell-surface PrP protein. Nature, 356, 577–582.

4. Richt, J.A., Kasinathan, P., Hamir, A.N., Castilla, J., Sathiyaseelan, T., Vargas, F., Sathiyaseelan, J., Wu, H., Matsushita, H., Koster, J., et al. (2007) Production of cattle lacking prion protein. Nat. Biotechnol., 25, 132–138.

5. Yu, G., Chen, J., Xu, Y., Zhu, C., Yu, H., Liu, S., Sha, H., Chen, J., Xu, X., Wu, Y., et al. (2009) Generation of goats lacking prion protein. Mol. Reprod. Dev., 76, 3.

6. Bremer, J., Baumann, F., Tiberi, C., Wessig, C., Fischer, H., Schwarz, P., Steele, A.D., Toyka, K.V., Nave, K.-A., Weis, J., et al. (2010) Axonal prion protein is required for peripheral myelin maintenance. Nat. Neurosci., 13, 310–318.

7. Benestad, S.L., Austbø, L., Tranulis, M.A., Espenes, A. and Olsaker, I. (2012) Healthy goats naturally devoid of prion protein. Vet. Res., 43, 87.

8. Minikel, E.V., Vallabh, S.M., Lek, M., Estrada, K., Samocha, K.E., Sathirapongsasuti, J.F., McLean, C.Y., Tung, J.Y., Yu, L.P.C., Gambetti, P., et al. (2016) Quantifying prion disease penetrance using large population control cohorts. Sci. Transl. Med., 8, 322ra9.

9. Minikel, E.V., Karczewski, K.J., Martin, H.C., Cummings, B.B., Whiffin, N., Rhodes, D., Alföldi, J., Trembath, R.C., van Heel, D.A., Daly, M.J., et al. (2020) Evaluating drug targets through human loss-of-function genetic variation. Nature, 581, 459–464.

10. Büeler, H., Raeber, A., Sailer, A., Fischer, M., Aguzzi, A. and Weissmann, C. (1994) High prion and PrPSc levels but delayed onset of disease in scrapie-inoculated mice heterozygous for a disrupted PrP gene. Mol. Med. Camb. Mass, 1, 19–30.

11. Büeler, H., Aguzzi, A., Sailer, A., Greiner, R.A., Autenried, P., Aguet, M. and Weissmann, C. (1993) Mice devoid of PrP are resistant to scrapie. Cell, 73, 1339–1347.

12. Sakaguchi, S., Katamine, S., Shigematsu, K., Nakatani, A., Moriuchi, R., Nishida, N., Kurokawa, K., Nakaoke, R., Sato, H. and Jishage, K. (1995) Accumulation of proteinase K-resistant prion protein (PrP) is restricted by the expression level of normal PrP in mice inoculated with a mouse-adapted strain of the Creutzfeldt-Jakob disease agent. J. Virol., 69, 7586–7592.

13. Fischer, M., Rülicke, T., Raeber, A., Sailer, A., Moser, M., Oesch, B., Brandner, S., Aguzzi, A. and Weissmann, C. (1996) Prion protein (PrP) with amino-proximal deletions restoring susceptibility of PrP knockout mice to scrapie. EMBO J., 15, 1255–1264.

14. Mallucci, G., Dickinson, A., Linehan, J., Klöhn, P.-C., Brandner, S. and Collinge, J. (2003) Depleting neuronal PrP in prion infection prevents disease and reverses spongiosis. Science, 302, 871–874.

15. Safar, J.G., DeArmond, S.J., Kociuba, K., Deering, C., Didorenko, S., Bouzamondo-Bernstein, E., Prusiner, S.B. and Tremblay, P. (2005) Prion clearance in bigenic mice. J. Gen. Virol., 86, 2913–2923.

16. Nazor Friberg, K., Hung, G., Wancewicz, E., Giles, K., Black, C., Freier, S., Bennett, F., Dearmond, S.J., Freyman, Y., Lessard, P., et al. (2012) Intracerebral Infusion of Antisense Oligonucleotides Into Prion-infected Mice. Mol. Ther. Nucleic Acids, 1, e9.

17. Raymond, G.J., Zhao, H.T., Race, B., Raymond, L.D., Williams, K., Swayze, E.E., Graffam, S., Le, J., Caron, T., Stathopoulos, J., et al. (2019) Antisense oligonucleotides extend survival of prion-infected mice. JCI Insight, 5.

18. Minikel, E.V., Zhao, H.T., Le, J., O’Moore, J., Pitstick, R., Graffam, S., Carlson, G.A., Kavanaugh, M.P., Kriz, J., Kim, J.B., et al. (2020) Prion protein lowering is a disease-modifying therapy across prion disease stages, strains and endpoints. Nucleic Acids Res., 10.1093/nar/gkaa616.

19. An, M., Davis, J.R., Levy, J.M., Serack, F.E., Harvey, J.W., Brauer, P.P., Pirtle, C.P., Berríos, K.N., Newby, G.A., Yeh, W.-H., et al. (2025) In vivo base editing extends lifespan of a humanized mouse model of prion disease. Nat. Med., 10.1038/s41591-024-03466-w.

20. Hay, M., Thomas, D.W., Craighead, J.L., Economides, C. and Rosenthal, J. (2014) Clinical development success rates for investigational drugs. Nat. Biotechnol., 32, 40–51.

21. Wong, C.H., Siah, K.W. and Lo, A.W. (2019) Estimation of clinical trial success rates and related parameters. Biostat. Oxf. Engl., 20, 273–286.

22. Thomas, D., Chancellor, D., Micklus, A., LaFever, S., Hay, M., Chaudhuri, S., Bowden, R. and Lo, A.W. (2021) Clinical Development Success Rates and Contributing Factors 2011–2020.

23. Minikel, E.V., Painter, J.L., Dong, C.C. and Nelson, M.R. (2024) Refining the impact of genetic evidence on clinical success. Nature, 10.1038/s41586-024-07316-0.

24. Sandberg, M.K., Al-Doujaily, H., Sharps, B., De Oliveira, M.W., Schmidt, C., Richard-Londt, A., Lyall, S., Linehan, J.M., Brandner, S., Wadsworth, J.D.F., et al. (2014) Prion neuropathology follows the accumulation of alternate prion protein isoforms after infective titre has peaked. Nat. Commun., 5, 4347.

25. Wu, H., Lima, W.F., Zhang, H., Fan, A., Sun, H. and Crooke, S.T. (2004) Determination of the role of the human RNase H1 in the pharmacology of DNA-like antisense drugs. J. Biol. Chem., 279, 17181–17189.

26. Tang, Q. and Khvorova, A. (2024) RNAi-based drug design: considerations and future directions. Nat. Rev. Drug Discov., 23, 341–364.

27. Dowdy, S.F. (2023) Endosomal escape of RNA therapeutics: How do we solve this rate-limiting problem? RNA N. Y. N, 29, 396–401.

28. Alterman, J.F., Godinho, B.M.D.C., Hassler, M.R., Ferguson, C.M., Echeverria, D., Sapp, E., Haraszti, R.A., Coles, A.H., Conroy, F., Miller, R., et al. (2019) A divalent siRNA chemical scaffold for potent and sustained modulation of gene expression throughout the central nervous system. Nat. Biotechnol., 37, 884–894.

29. Ferguson, C.M., Hildebrand, S., Godinho, B.M.D.C., Buchwald, J., Echeverria, D., Coles, A., Grigorenko, A., Vangjeli, L., Sousa, J., McHugh, N., et al. (2024) Silencing Apoe with divalent-siRNAs improves amyloid burden and activates immune response pathways in Alzheimer’s disease. Alzheimers Dement. J. Alzheimers Assoc., 20, 2632–2652.

30. Andreone, B.J., Lin, J., Tocci, J., Rook, M., Omer, A., Carito, L.M., Yang, C., Zhoba, H., DeJesus, C., Traore, M., et al. (2025) Durable suppression of seizures in a preclinical model of KCNT1 genetic epilepsy with divalent small interfering RNA. Epilepsia, 66, 1677–1690.

31. Weiss, A., Gilbert, J.W., Rivera Flores, I.V., Belgrad, J., Ferguson, C., Dogan, E.O., Wightman, N., Mocarski, K., Echeverria, D., Harkins, A.L., et al. (2025) RNAi-mediated silencing of SOD1 profoundly extends survival and functional outcomes in ALS mice. Mol. Ther. J. Am. Soc. Gene Ther., 33, 3917–3938.

32. Crooke, S.T., Vickers, T.A. and Liang, X.-H. (2020) Phosphorothioate modified oligonucleotide-protein interactions. Nucleic Acids Res., 48, 5235–5253.

33. Shen, W., De Hoyos, C.L., Migawa, M.T., Vickers, T.A., Sun, H., Low, A., Bell, T.A., Rahdar, M., Mukhopadhyay, S., Hart, C.E., et al. (2019) Chemical modification of PS-ASO therapeutics reduces cellular protein-binding and improves the therapeutic index. Nat. Biotechnol., 37, 640–650.

34. Yamada, K., Hariharan, V.N., Caiazzi, J., Miller, R., Ferguson, C.M., Sapp, E., Fakih, H.H., Tang, Q., Yamada, N., Furgal, R.C., et al. (2024) Enhancing siRNA efficacy in vivo with extended nucleic acid backbones. Nat. Biotechnol., 10.1038/s41587-024-02336-7.

35. De, N., Young, L., Lau, P.-W., Meisner, N.-C., Morrissey, D.V. and MacRae, I.J. (2013) Highly complementary target RNAs promote release of guide RNAs from human Argonaute2. Mol. Cell, 50, 344–355.

36. Sheu-Gruttadauria, J., Pawlica, P., Klum, S.M., Wang, S., Yario, T.A., Schirle Oakdale, N.T., Steitz, J.A. and MacRae, I.J. (2019) Structural Basis for Target-Directed MicroRNA Degradation. Mol. Cell, 75, 1243–1255.e7.

37. Tang, Q., Fakih, H.H., Zain Ui Abideen, M., Hildebrand, S.R., Afshari, K., Gross, K.Y., Sousa, J., Maebius, A.S., Bartholdy, C., Søgaard, P.P., et al. (2023) Rational design of a JAK1-selective siRNA inhibitor for the modulation of autoimmunity in the skin. Nat. Commun., 14, 7099.

38. Shmushkovich, T., Monopoli, K.R., Homsy, D., Leyfer, D., Betancur-Boissel, M., Khvorova, A. and Wolfson, A.D. (2018) Functional features defining the efficacy of cholesterol-conjugated, self-deliverable, chemically modified siRNAs. Nucleic Acids Res., 46, 10905–10916.

39. Ballmer, B.A., Moos, R., Liberali, P., Pelkmans, L., Hornemann, S. and Aguzzi, A. (2017) Modifiers of prion protein biogenesis and recycling identified by a highly parallel endocytosis kinetics assay. J. Biol. Chem., 292, 8356–8368.

40. Gentile, J.E., Corridon, T.L., Mortberg, M.A., D’Souza, E.N., Whiffin, N., Minikel, E.V. and Vallabh, S.M. (2024) Modulation of prion protein expression through cryptic splice site manipulation. J. Biol. Chem., 300, 107560.

41. Janas, M.M., Schlegel, M.K., Harbison, C.E., Yilmaz, V.O., Jiang, Y., Parmar, R., Zlatev, I., Castoreno, A., Xu, H., Shulga-Morskaya, S., et al. (2018) Selection of GalNAc-conjugated siRNAs with limited off-target-driven rat hepatotoxicity. Nat. Commun., 9, 723.

42. Nuvolone, M., Hermann, M., Sorce, S., Russo, G., Tiberi, C., Schwarz, P., Minikel, E., Sanoudou, D., Pelczar, P. and Aguzzi, A. (2016) Strictly co-isogenic C57BL/6J-Prnp-/-mice: A rigorous resource for prion science. J. Exp. Med., 213, 313–327.

43. de Vree, P.J.P., de Wit, E., Yilmaz, M., van de Heijning, M., Klous, P., Verstegen, M.J.A.M., Wan, Y., Teunissen, H., Krijger, P.H.L., Geeven, G., et al. (2014) Targeted sequencing by proximity ligation for comprehensive variant detection and local haplotyping. Nat. Biotechnol., 32, 1019–1025.

44. Vallabh, S.M., Minikel, E.V., Williams, V.J., Carlyle, B.C., McManus, A.J., Wennick, C.D., Bolling, A., Trombetta, B.A., Urick, D., Nobuhara, C.K., et al. (2020) Cerebrospinal fluid and plasma biomarkers in individuals at risk for genetic prion disease. BMC Med., 18, 140.

45. Miller, R., Paquette, J., Barker, A., Sapp, E., McHugh, N., Bramato, B., Yamada, N., Alterman, J., Echeveria, D., Yamada, K., et al. (2024) Preventing acute neurotoxicity of CNS therapeutic oligonucleotides with the addition of Ca2+ and Mg2+ in the formulation. Mol. Ther. Nucleic Acids, 35, 102359.

46. Moazami, M.P., Rembetsy-Brown, J.M., Sarli, S.L., McEachern, H.R., Wang, F., Ohara, M., Wagh, A., Kelly, K., Krishnamurthy, P.M., Weiss, A., et al. (2024) Quantifying and mitigating motor phenotypes induced by antisense oligonucleotides in the central nervous system. Mol. Ther. J. Am. Soc. Gene Ther., 10.1016/j.ymthe.2024.10.024.

47. Hernandez, M.B., Mazur, C., Chen, H., Fradkin, L., Searcy, J., Burel, S., Kelly, M., Bruening, D., O’Rourke, J.G., Cai, Y., et al. (2025) Transient Acute Neuronal Activation Response Caused by High Concentrations of Oligonucleotides in the Cerebral Spinal Fluid. bioRxiv, 10.1101/2025.02.13.638138.

48. Jones, S. Avertin Solution for Mouse Anesthesia.

49. Corridon, T.L., O’Moore, J., Lian, Y., Laversenne, V., Noble, B., Kamath, N.G., Serack, F.E., Shaikh, A.B., Erickson, B., Braun, C., et al. (2024) PrP turnover in vivo and the time to effect of prion disease therapeutics. bioRxiv, 10.1101/2024.11.12.623215.

50. Chandler, R.L. (1961) Encephalopathy in mice produced by inoculation with scrapie brain material. Lancet Lond. Engl., 1, 1378–1379.

51. Mortberg, M.A., Zhao, H.T., Reidenbach, A.G., Gentile, J.E., Kuhn, E., O’Moore, J., Dooley, P.M., Connors, T.R., Mazur, C., Allen, S.W., et al. (2022) Regional variability and genotypic and pharmacodynamic effects on PrP concentration in the CNS. JCI Insight, 7, e156532.

52. Reidenbach, A.G., Mesleh, M.F., Casalena, D., Vallabh, S.M., Dahlin, J.L., Leed, A.J., Chan, A.I., Usanov, D.L., Yehl, J.B., Lemke, C.T., et al. (2020) Multimodal small-molecule screening for human prion protein binders. J. Biol. Chem., 295, 13516–13531.

53. Godinho, B.M.D.C., Gilbert, J.W., Haraszti, R.A., Coles, A.H., Biscans, A., Roux, L., Nikan, M., Echeverria, D., Hassler, M. and Khvorova, A. (2017) Pharmacokinetic Profiling of Conjugated Therapeutic Oligonucleotides: A High-Throughput Method Based Upon Serial Blood Microsampling Coupled to Peptide Nucleic Acid Hybridization Assay. Nucleic Acid Ther., 27, 323–334.

54. Gao, X., Diep, J.K., Norris, D.A., Yu, R.Z. and Geary, R.S. (2023) Predicting the pharmacokinetics and pharmacodynamics of antisense oligonucleotides: an overview of various approaches and opportunities for PBPK/PD modelling. Expert Opin. Drug Metab. Toxicol., 19, 979–990.

55. Khvorova, A. and Kennedy, Z. (2021) Oligonucleotides for prnp modulation.

56. Tang, T., Li, L., Tang, J., Li, Y., Lin, W.Y., Martin, F., Grant, D., Solloway, M., Parker, L., Ye, W., et al. (2010) A mouse knockout library for secreted and transmembrane proteins. Nat. Biotechnol., 28, 749–755.

57. Dickinson, M.E., Flenniken, A.M., Ji, X., Teboul, L., Wong, M.D., White, J.K., Meehan, T.F., Weninger, W.J., Westerberg, H., Adissu, H., et al. (2016) High-throughput discovery of novel developmental phenotypes. Nature, 537, 508–514.

58. Frmd6 | FERM domain containing 6 mouse gene | IMPC Int. Mouse Phenotyping Consort. IMPC.

59. Schirle, N.T., Sheu-Gruttadauria, J. and MacRae, I.J. (2014) Structural basis for microRNA targeting. Science, 346, 608–613.

60. Becker, W.R., Ober-Reynolds, B., Jouravleva, K., Jolly, S.M., Zamore, P.D. and Greenleaf, W.J. (2019) High-Throughput Analysis Reveals Rules for Target RNA Binding and Cleavage by AGO2. Mol. Cell, 75, 741–755.e11.

61. Wang, P.Y. and Bartel, D.P. (2024) The guide-RNA sequence dictates the slicing kinetics and conformational dynamics of the Argonaute silencing complex. Mol. Cell, 84, 2918–2934.e11.

62. Freier, S.M., Bui, H.-H. and Zhao, H. (2020) Compounds and methods for reducing prion expression.

63. Emami, A., Tepper, J., Short, B., Yaksh, T.L., Bendele, A.M., Ramani, T., Cisternas, A.F., Chang, J.H. and Mellon, R.D. (2018) Toxicology Evaluation of Drugs Administered via Uncommon Routes: Intranasal, Intraocular, Intrathecal/Intraspinal, and Intra-Articular. Int. J. Toxicol., 37, 4–27.

64. Stern, S., Wange, R.L. and Rogers, H. (2024) An Evaluation of First-in-Human Studies for RNA Oligonucleotides. Nucleic Acid Ther., 10.1089/nat.2024.0036.

65. Miller, T.M., Cudkowicz, M.E., Genge, A., Shaw, P.J., Sobue, G., Bucelli, R.C., Chiò, A., Van Damme, P., Ludolph, A.C., Glass, J.D., et al. (2022) Trial of Antisense Oligonucleotide Tofersen for SOD1 ALS. N. Engl. J. Med., 387, 1099–1110.

66. Jafar-Nejad, P., Powers, B., Soriano, A., Zhao, H., Norris, D.A., Matson, J., DeBrosse-Serra, B., Watson, J., Narayanan, P., Chun, S.J., et al. (2021) The atlas of RNase H antisense oligonucleotide distribution and activity in the CNS of rodents and non-human primates following central administration. Nucleic Acids Res., 49, 657–673.

67. Frei, J.A., Gentile, J.E., Lian, Y., Mortberg, M.A., Capitanio, J., Jafar-Nejad, P., Vallabh, S.M., Zhao, H.T. and Minikel, E.V. (2025) Cell Type Distribution of Intrathecal Antisense Oligonucleotide Activity in Deep Brain Regions of Non-Human Primates. Nucleic Acid Ther., 10.1177/21593337251371594.

68. McDonough, S. Widespread and Durable Knockdown of HTT in Non-Human Primate Brain by a Novel Oligonucleotide Modality. In. Palm Springs, CA.

69. Ferguson, C.M., Godinho, B.M., Alterman, J.F., Coles, A.H., Hassler, M., Echeverria, D., Gilbert, J.W., Knox, E.G., Caiazzi, J., Haraszti, R.A., et al. (2021) Comparative route of administration studies using therapeutic siRNAs show widespread gene modulation in Dorset sheep. JCI Insight, 6, e152203.

70. Brandel, J.-P., Vlaicu, M.B., Culeux, A., Belondrade, M., Bougard, D., Grznarova, K., Denouel, A., Plu, I., Bouaziz-Amar, E., Seilhean, D., et al. (2020) Variant Creutzfeldt-Jakob Disease Diagnosed 7.5 Years after Occupational Exposure. N. Engl. J. Med., 383, 83–85.

71. Casassus, B. (2021) France halts prion research amid safety concerns. Science, 373, 475– 476.

72. Chou, S.-W., Mortberg, M.A., Marlen, K., Ojala, D.S., Parman, T., Howard, M., DeSouza-Lenz, K., Lian, Y., Mehrabian, M., Tiffany, M., et al. (2025) Zinc Finger Repressors mediate widespread PRNP lowering in the nonhuman primate brain and profoundly extend survival in prion disease mice. 10.1101/2025.03.05.636713.

73. Wadman, M. (2024) Foiling deadly prions. Science, 383.

