## Supplement for "Divalent siRNA for prion disease"

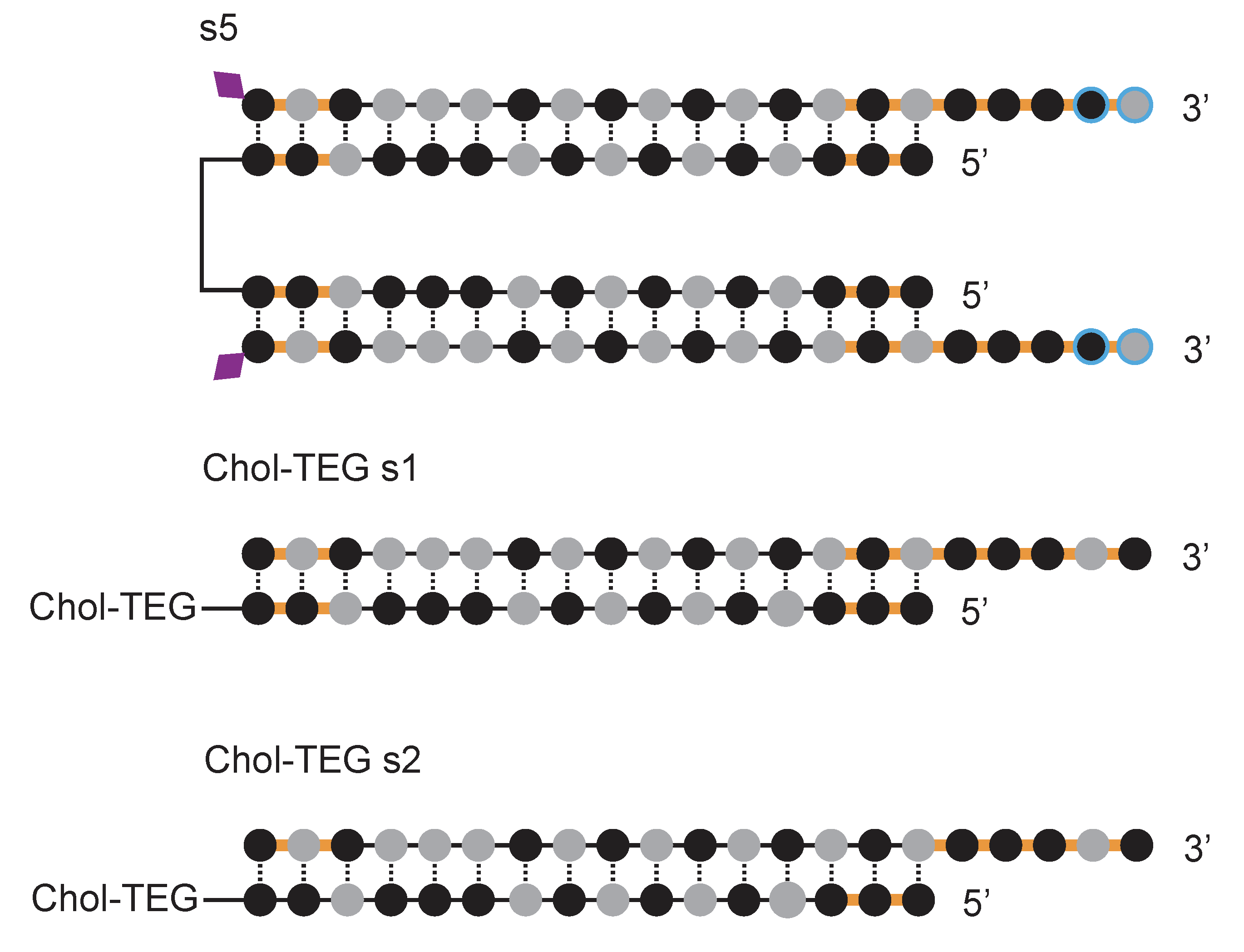

***Figure S1. Additional chemical scaffolds.*** *Scaffold s5 is utilized in Figure S9. The Chol-TEG scaffolds were used in cellular screening assays.*

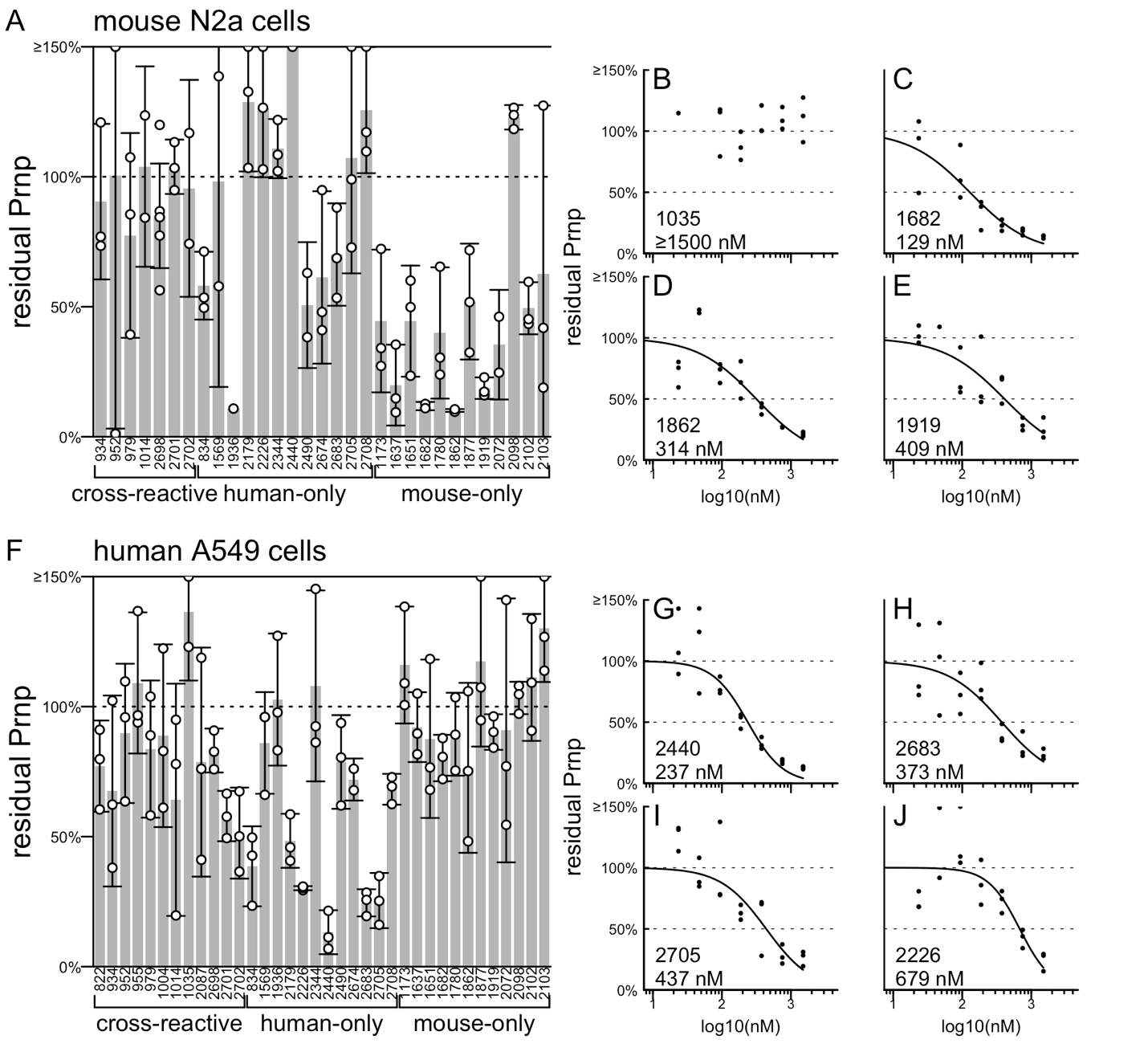

***Figure S2. Initial screening by bDNA assay in cell culture.*** *These screens used scaffold Chol-TEG s1 (Figure S1).* ***A)*** *Mouse N2a cells, each point is one well, triplicate wells are analyzed for each compound, rectangular bars are means, error bars are 95% confidence intervals. Readout is Prnp expression normalized to Hprt, further normalized to the mean of predicted non-targeting (human-only) compounds.* ***B-E)*** *IC_50_ determination for 4 compounds selected from mouse N2a cell screen. Each point is one well, triplicate wells are tested at each dose level, curves are 4-parameter log-logistic dose-response curves fit using the drc package in R.* ***F)*** *As in (A) but for human A549 cells.* ***G-J)*** *As in (B-E) but for top human sequences in human A549 cells.*

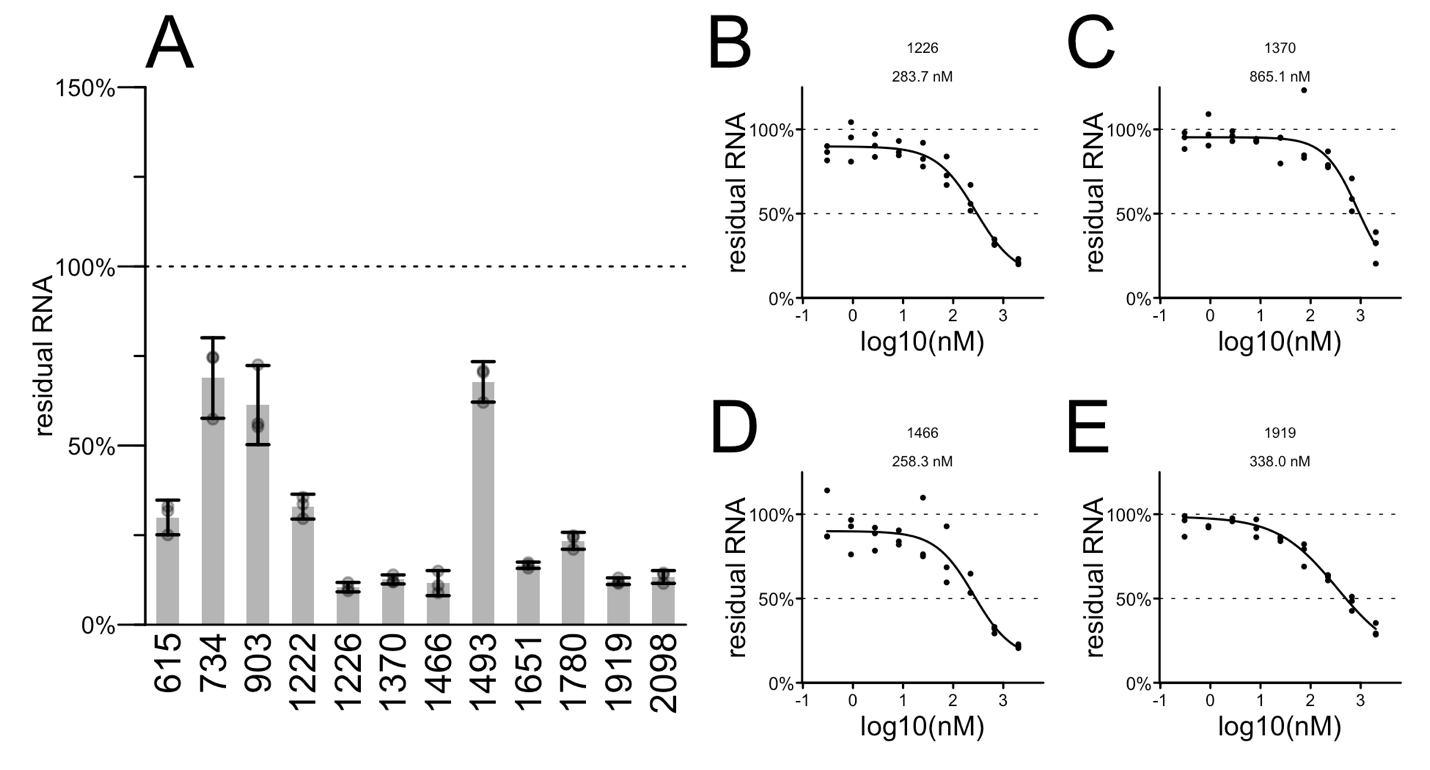
***Figure S3. Further screening of potential siRNA tool compounds against mouse Prnp in N2a cells.*** *This screen used scaffold Chol-TEG s2 (Figure S1).* ***A)*** *Each point is one well, triplicate wells are analyzed for each compound, readout is RT-qPCR with Prnp Ct values normalized to Tbp, then each point is normalized to the mean of untreated wells. Rectangular bars are means, error bars are 95% confidence intervals.* ***B-E)*** *IC_50_ determination for top compounds. Each point is one well, triplicate wells are tested at each dose level, curves are 4-parameter log-logistic dose-response curves fit using the drc package in R.*

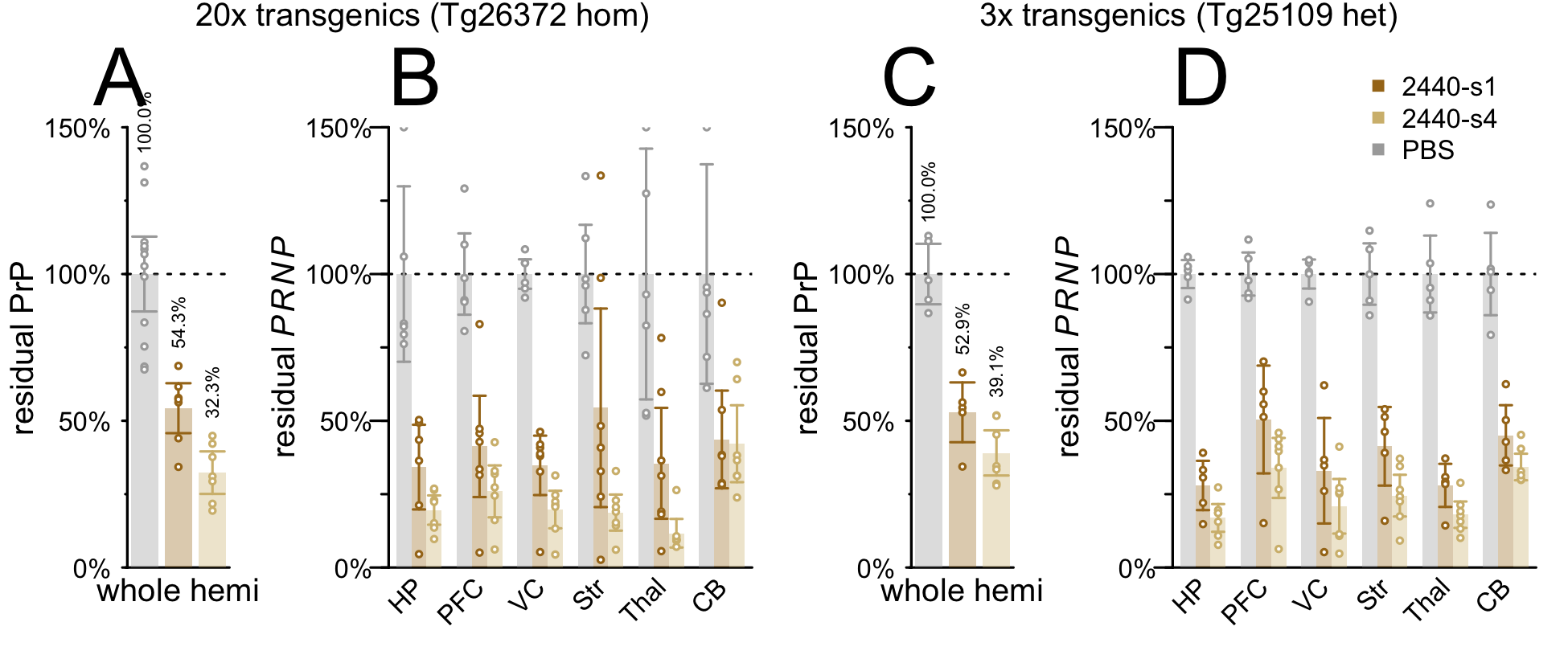

***Figure S4. Potency of tool compound 2440 in high or low copy number HuPrP transgenic mice. A-B)*** *2440-s1 and 2440-s4 tested at 348 µg in Tg26372 homozygous mice with 20 copies of PRNP and 5.4- fold PrP expression.* ***A)*** *Whole hemisphere PrP ELISA readout. These data are reproduced from Figure 4C for convenience of comparison.* ***B)*** *Regional RT-qPCR readout.* ***C-D)*** *2440-s1 and 2440-s4 tested at 348 µg in Tg25109 heterozygous mice with 3 copies of PRNP and 1.1-fold PrP expression.* ***C)*** *Whole hemisphere PrP ELISA readout.* ***D)*** *Regional RT-qPCR readout. For all panels, each point is one animal, rectangular bars are means, error bars are 95% confidence intervals.*

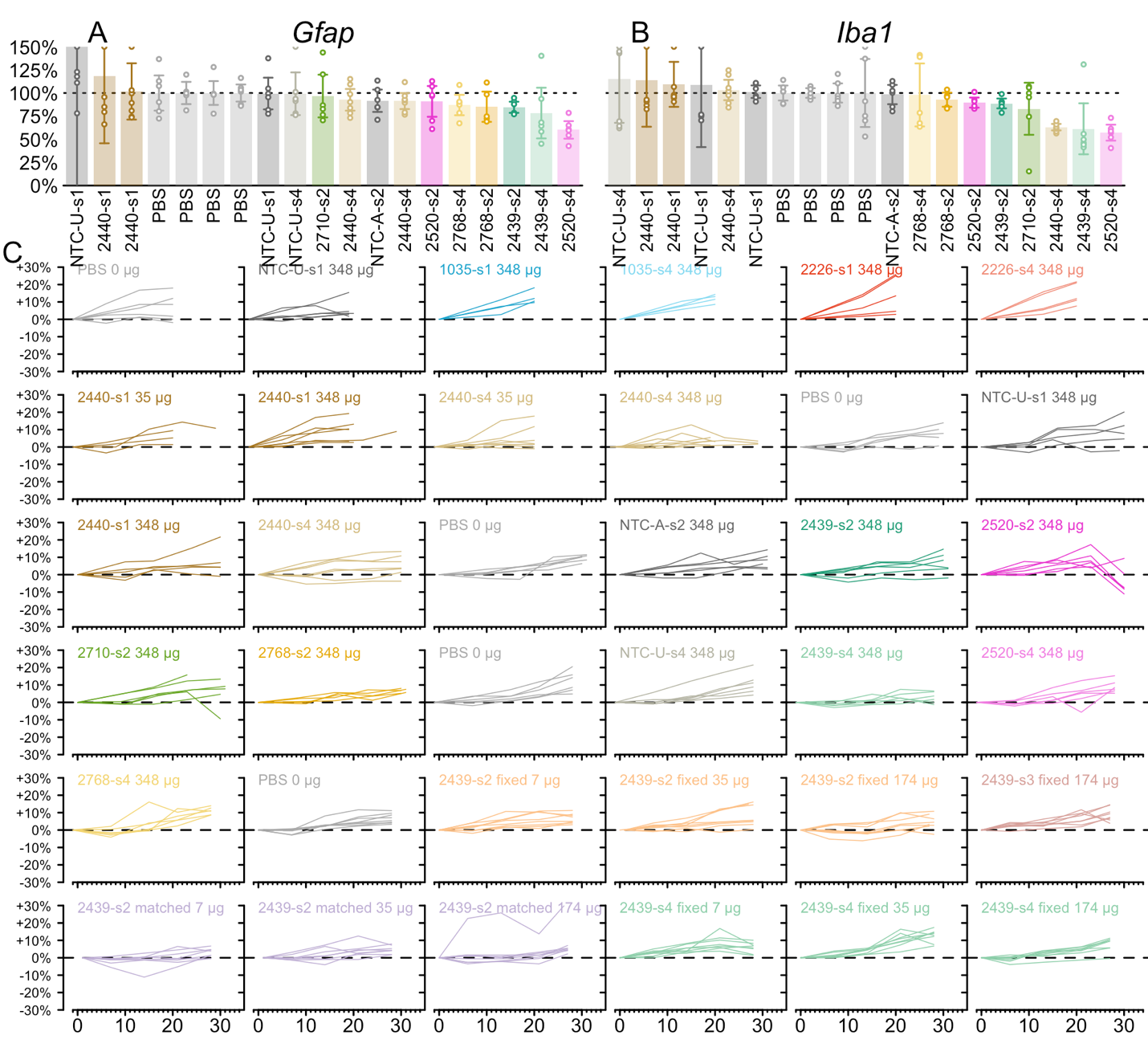

***Figure S5. Tolerability metrics for divalent siRNAs in short-term target engagement studies. A)*** *Gfap and* ***B)*** *Iba1 by RT-qPCR in visual cortex for all studies in Figure 4 and Figure 5, normalized to the mean of PBS controls within each study. Each point is one animal. Rectangular bars are means, error bars are 95% confidence intervals. Compounds are sorted along the x axis by rank of mean expression of each inflammatory marker. Each cohort is normalized to the PBS control within its own experiment, and each PBS control group is displayed separately.* ***C)*** *Individual weight gain trajectories for every animal in Figure 4 and 5, normalized to individual baseline. Weight change from individual baseline as a percent (y axis) versus days post-dose (x axis). The PBS control group from each experiment is displayed separately.*

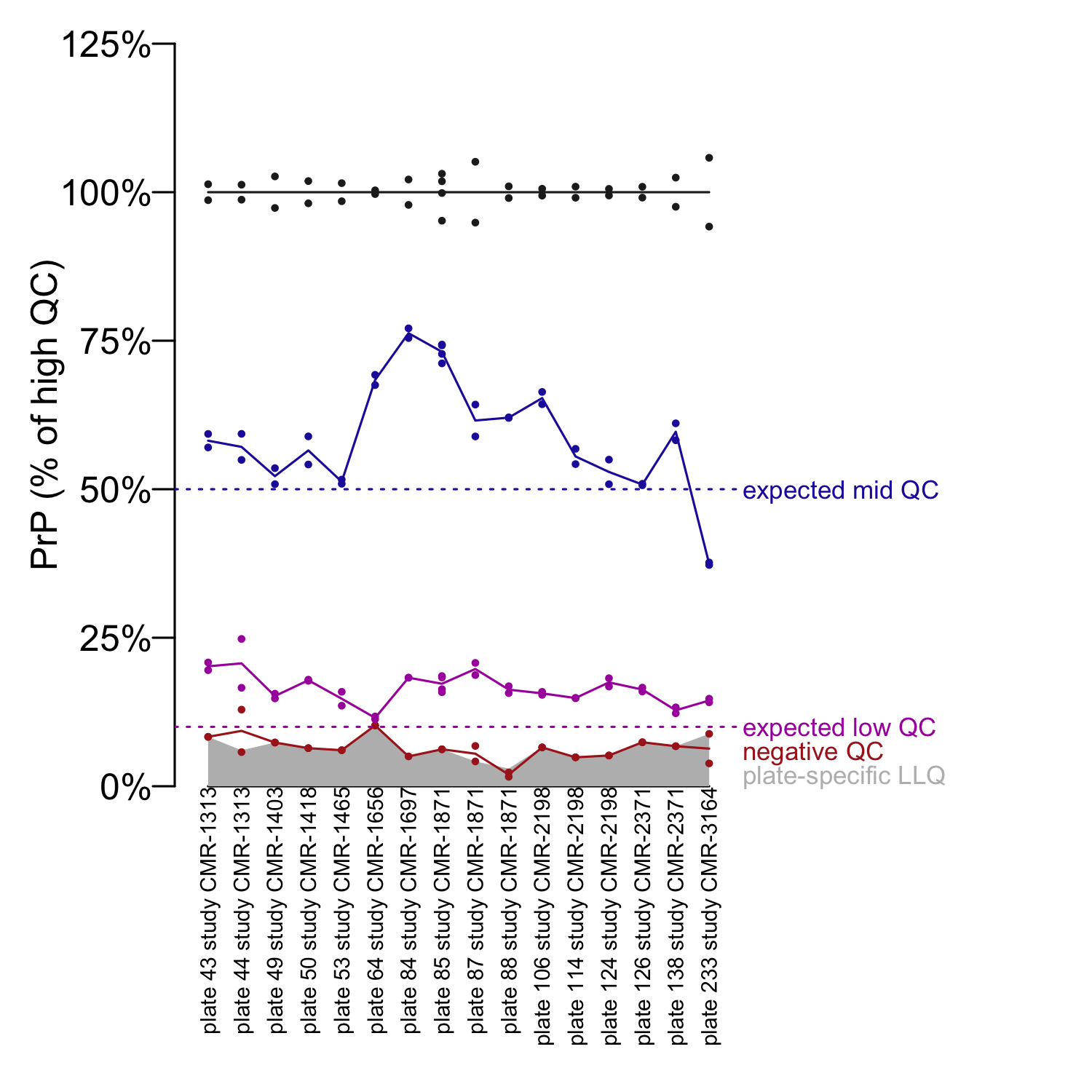
***Figure S6. Quality control metrics for PrP ELISA plates.*** *Every plate includes, in duplicate, the same high QC (WT mouse brain, black), mid QC (het KO mouse brain, blue), low QC (contrived sample of 90% KO brain homogenate spiked with 10% WT brain homogenate to simulate 10% residual PrP, magenta) and negative QC (KO brain, maroon). The LLQ is 0.05 ng/mL, and QCs are run at a final 1:200 dilution so that 10 ng PrP per g of wet brain tissue is the lower limit of quantification for these samples. In this plot, each point is one replicate of a QC, and its PrP concentration is normalized to the mean of high QCs. Readings from consecutive plates are connected by lines. The low QC, designed to simulate 10% residual PrP, read out at a mean of 17% of high QC across all plates shown here, slightly higher than the 14% found in validation (48). The source of this floor effect is unknown; one possibility is promiscuous binding to non-PrP proteins.*

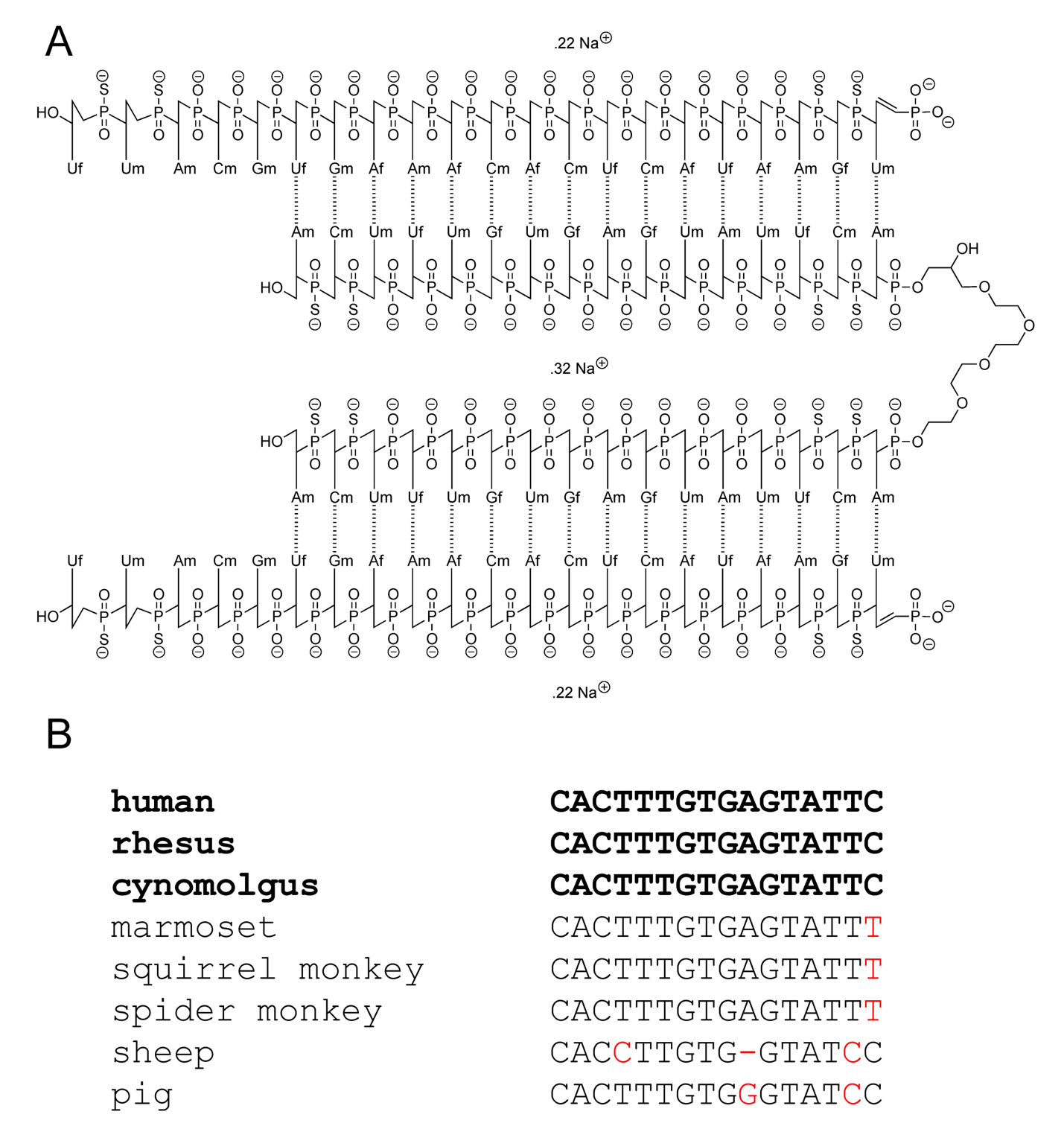
***Figure S7. Identity and cross-species reactivity analysis for sequence 2439. A)*** *Chemical structure of 2439-s4. f = 2′ Fluoro, m = 2′-O-methyl.* ***B)*** *Multiple species alignment of genomic sequences complementing bases 2-17 of the antisense strand.* *Fully matched sequences are shown in bold black. For imperfectly matched species, matched bases in black, indels or mismatches in red. No alignments were found for mouse, rat, Syrian hamster, or dog.*

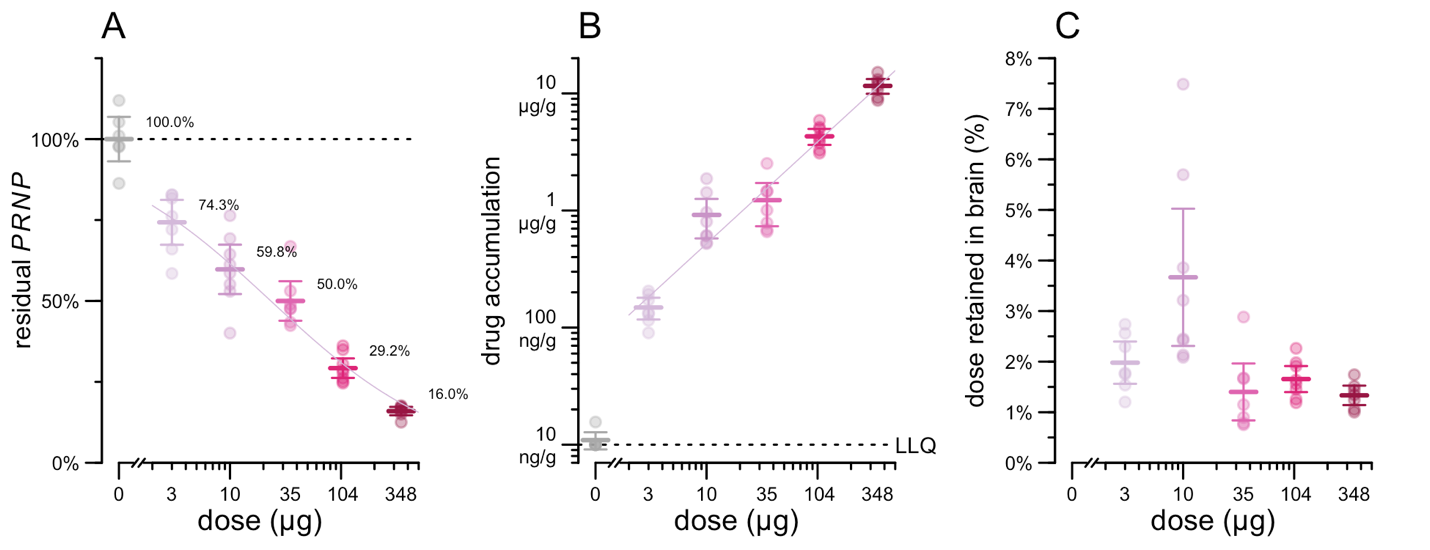

***Figure S8. Additional analyses of pharmacodynamic/pharmacokinetic data in humanized mice. A)*** *Administered dose (x axis) vs. residual PRNP mRNA in whole brain hemisphere (y axis).* ***B)*** *Administered dose (x axis) vs. drug accumulation in whole brain hemisphere (y axis).* ***C)*** *Administered dose (x axis) vs. percentage of administered dose retained in brain (assuming a 0.4 g brain times the measured tissue concentration in µg/g; y axis).*

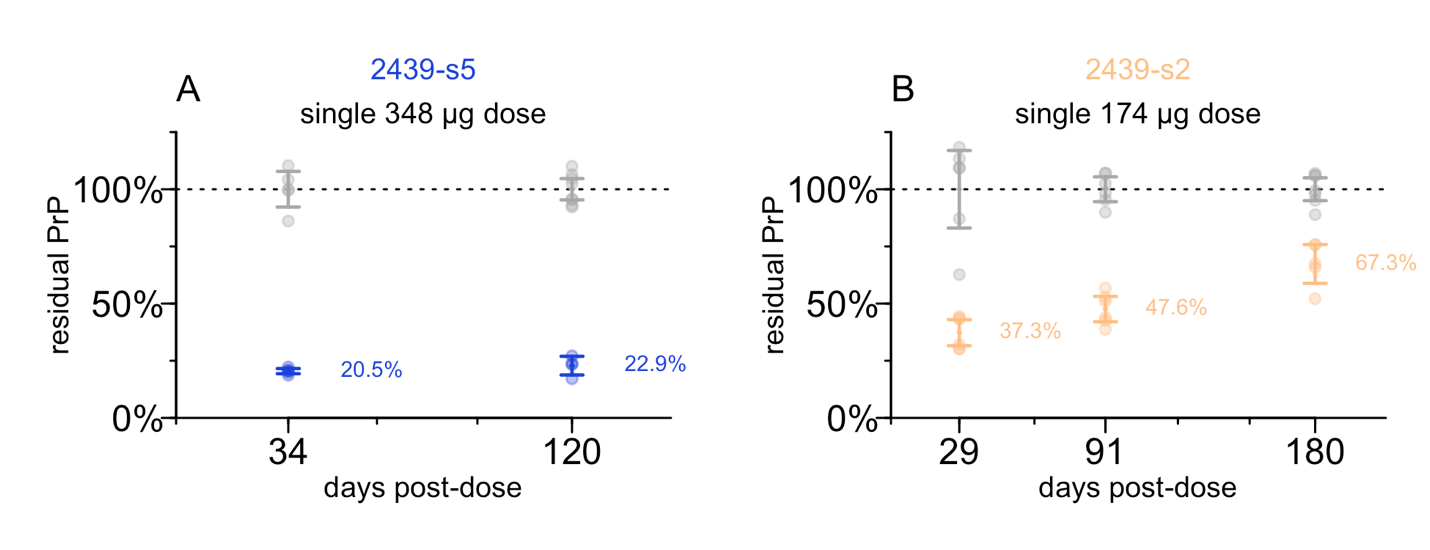

***Figure S9. Durability studies for 2439 in other scaffolds. A)*** *2439-s5 (see Figure S1 for scaffold description) at 348 µg.* ***B)*** *2439-s2 at 174 µg. The same control animals from Figure 6 are reproduced here for convenience of comparison.*

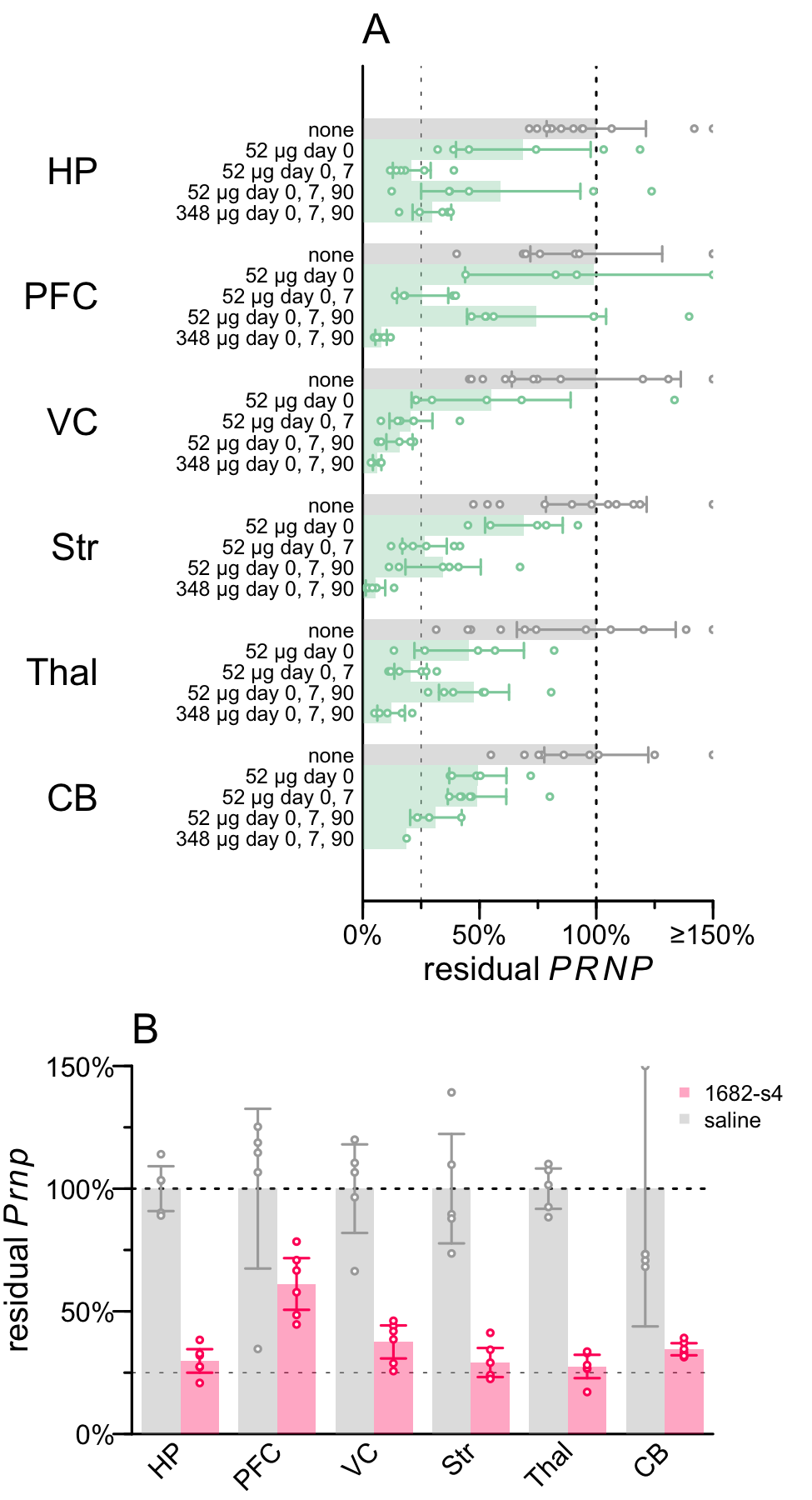

***Figure S10. Regional qPCR analysis for repeat dose 2439-s4 versus 1682-s4.*** *Regions: hippocampus (HP), prefrontal cortex (PFC), visual cortex (VC), striatum (Str), thalamus (Thal), and cerebellum (CB).* ***A)*** *Regional PRNP RT-qPCR analysis for repeat dose 2439-s4 study in the same Tg26372 animals shown in Figure 6D.* ***B)*** *Regional Prnp RT-qPCR for 1682-s4 from the same animals shown in Figure 2A.*

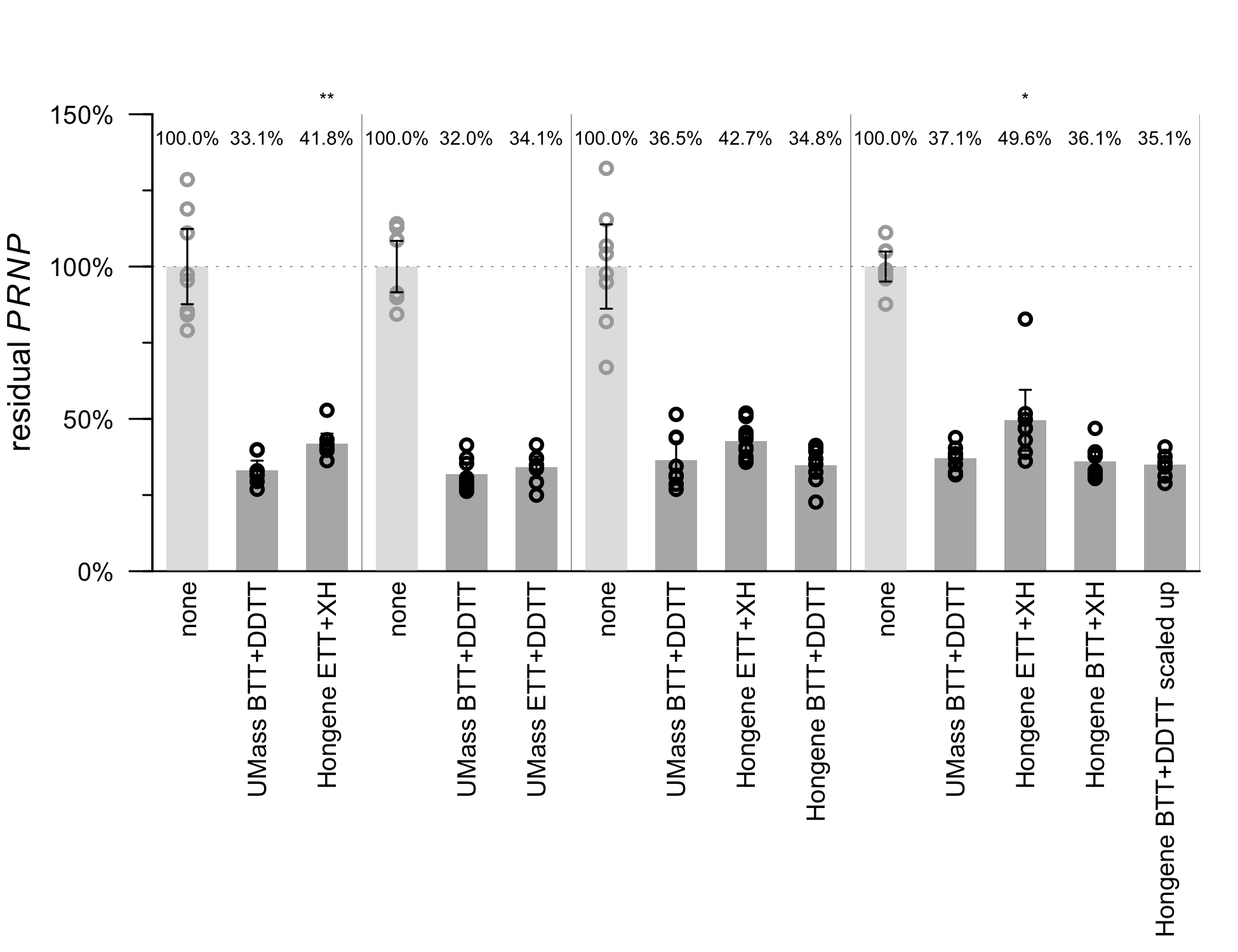

***Figure S11. Potency assessment for batches of 2439-s4 manufactured by different processes.*** *Batches were manufactured by UMass or Hongene using different activators (BTT or ETT) and sulfurization reagents (DDTT or XH). After the different combinations of reagents were tested, the Hongene BTT+DDTT process was scaled up (rightmost bar) in preparation for GMP synthesis, but all results shown in this figure are for non-GMP material. Each batch was injected into N=8-10 Tg26372 mice at a 139 µg dose level and whole hemispheres were analyzed by PRNP mRNA RT-qPCR at 7 days post-dose. *P < 0.05, **P < 0.01, for T test comparison to the UMass BTT+DDTT batch used as a reference.*

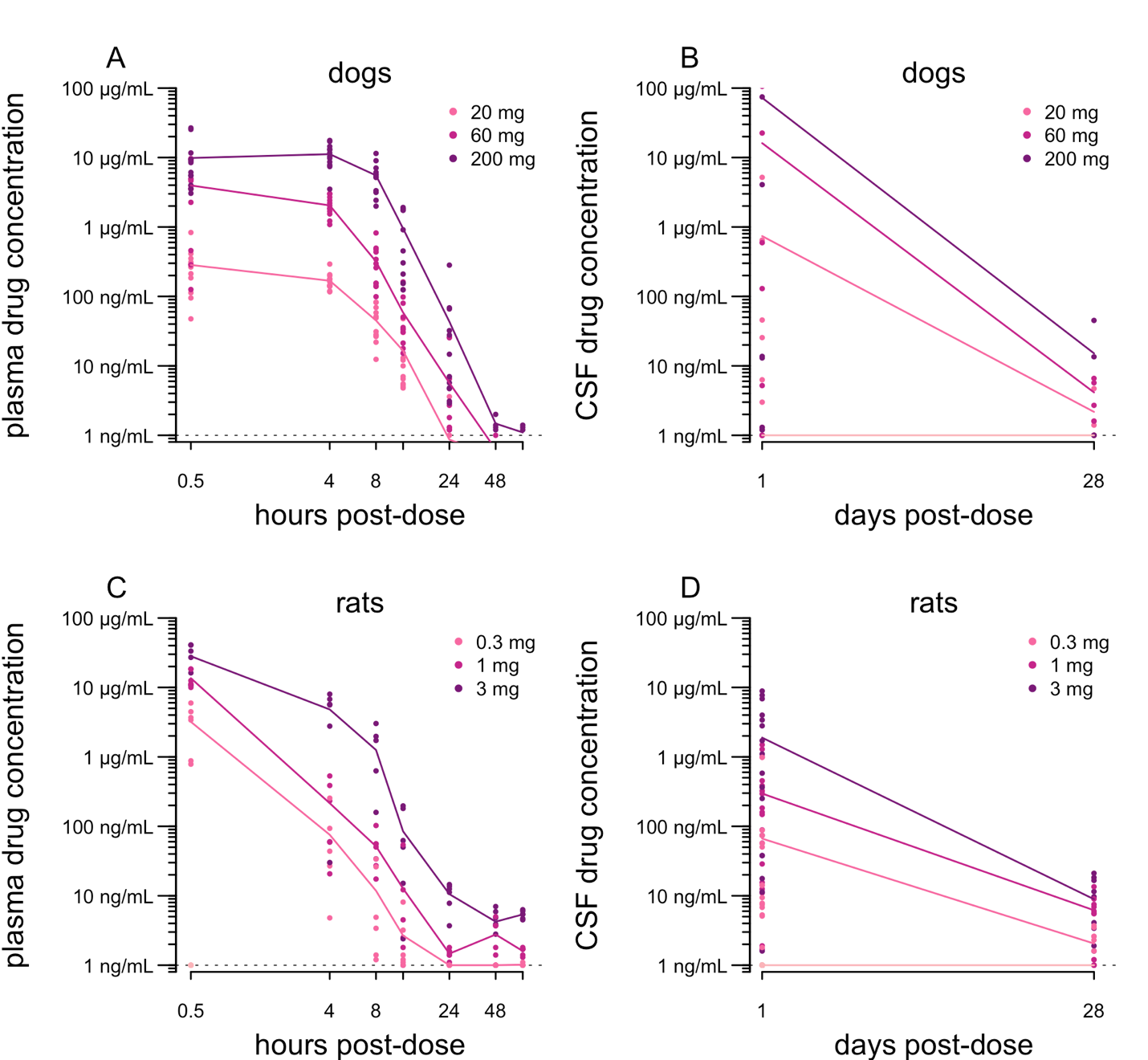

***Figure S12. CSF and plasma biodistribution analyses from GLP toxicology studies. A)*** *Dog plasma. N = 12 dogs per dose level per timepoint for 0.5 – 24 hours, N = 4 per timepoint for 48-72 hours.* ***B)*** *Dog CSF. N = 8 dogs per dose level at 1 day, N = 4 at 28 day.* ***C)*** *Rat plasma. N = 6 rats per timepoint.* ***D)*** *Rat CSF.* *N = 20 rats per dose level at 1 day, N=10 at 28 day. In all panels, line segments connect the means at each timepoint for each dose level.*
